# SHARP: Generating Synthesizable Molecules via Fragment-based Hierarchical Action-space Reinforcement Learning for Pareto Optimization

**DOI:** 10.1101/2025.07.18.665529

**Authors:** Jeonghyeon Kim, Seongok Ryu, Hahnbeom Park, Chaok Seok

## Abstract

Designing drug-like molecules that satisfy multiple objectives—such as high binding affinity, synthesizability, and drug-likeness—poses a complex global optimization problem over an astronomically large chemical space. Existing deep learning-based molecular generative models often treat this task as distribution modeling, relying on atom-level autoregressive actions with less consideration of explicit optimization feedback. Consequently, they frequently generate invalid structures, converge to local optima, or produce synthetically infeasible candidates. Here, we introduce SHARP (Synthesizable Hierarchical Action-space Reinforcement learning for Pareto optimization), a molecular generator that addresses these limitations via a fragment-based hierarchical action space and reinforcement learning. SHARP ensures synthetic accessibility by applying action masks guided by a pretrained Synthesizability Estimation Model (SEM). The reinforcement learning (RL) policy is trained using a composite reward function integrating docking scores, pharmacophore matching, and solvent accessibility to generate functionally relevant and experimentally tractable molecules. Furthermore, across four lead optimization tasks—fragment growing, linker design, scaffold hopping, and sidechain decoration—on a diverse receptor set, SHARP consistently outperforms prior methods in producing molecules at high affinity and synthesizability. These results demonstrate that reinforcement learning with a chemically intuitive action space design can be an effective solution to the optimization challenges in AI-driven drug discovery, offering a robust framework for rational molecular design in structure-based applications.

## Introduction

Designing novel drug-like molecules is fundamentally a multi-objective global optimization challenge. The task requires navigating a chemical space of astronomical size — estimated at over 10^60^ molecules^1^—to discover candidates that reside on the optimal Pareto front of synthesizability, binding affinity, and drug-likeness. Achieving this balance is critical because high potency is of little value without a viable synthetic route, and a readily synthesized molecule is irrelevant if it fails to bind the target. However, the majority of deep generative models are not structured to solve this optimization problem. Instead of explicitly seeking Pareto-optimal solutions, they are often conditioned on receptor structures and trained to replicate known chemical motifs, lacking the direct feedback from oracle functions — such as binding affinity and synthesizability scores — needed to guide the search toward superior, novel candidates.^2–4^

With advances in deep learning, numerous molecular generative models have emerged. Initially, ligand-based models generated molecules using only 2D graphs or SMILES.^5,6^ The advent of high-accuracy protein structure prediction methods such as AlphaFold2^7^ and RoseTTAFold^8^ enabled the integration of 3D protein-ligand interactions. Models such as LiGAN^9^ used CNNs, while the development of SE(3)-equivariant GNNs^10^ led to more sophisticated approaches like GraphBP,^11^ Pocket2Mol,^12^ or diffusion-based generative models such as DiffSBDD,^13^ DiffBP,^14^ and TargetDiff.^15^ While these models incorporate valuable structural information, their core design remains less suitable for molecular property optimization. They primarily learn to model a data distribution rather than to perform an explicit, objective-driven search for molecules that optimally balance synthesizability and binding efficacy.

We note that reliance on distribution modeling can lead to significant limitations. Such models are trained to replicate the empirical distribution of known ligand-protein complexes, which provides insufficient guidance for discovering molecules having both novelty and optimal drug properties. Lacking an explicit mechanism to maximize objective functions, these methods are constrained to exploring regions of the chemical space proximal to the training data. Consequently, they tend to generate molecules with limited structural diversity, often confined to known ligand configurations and scaffolds. This can pose a significant limitation in real-world drug design applications, such as discovering novel molecules or targeting previously unexplored receptors.

Two common methodological choices in previous methods can exacerbate the issues above. First, the use of an atom-based generation scheme frames the optimization within a dense and rugged search landscape. This high-dimensional, discrete space increases the probability that the algorithm will terminate at local minima far from the global optimum. Furthermore, an atomistic representation makes it exceedingly difficult to encode and optimize for synthesizability, as this property is governed by the relationships between larger, chemically relevant functional groups. Second, the prevalent autoregressive approach, where atoms are placed sequentially, is prone to error propagation. A chemically invalid placement in the initial steps can lock the generation process onto a path toward a less likely or invalid final molecule, as there is no mechanism to revise early decisions based on the context of the complete structure.

In this work, we introduce SHARP (Synthesizable Hierarchical Action-space Reinforcement learning for Pareto optimization), a framework that formulates molecular generation as a multi-objective global optimization problem. The unique aspect of SHARP is, first, its integrative RL components that can smoothly drive the search towards Pareto optimality, and second, its chemically intuitive action space, which allows the autoregressive generation process not only to grow but also to revise and refine molecular structures. In the following sections, the general concept of the SHARP model is described first. We then present detailed analyses of the Pareto optimization results, including methodological contributions to optimization performance, the generalizability of the method, and case study examples.

## Results and discussion

### Overview of the SHARP workflow

We first describe our model architecture, with an overview provided in Figure 1. SHARP follows a general reinforcement learning scheme, in which the action space search is guided by a reward function trained to effectively enhance binding affinity as well as other properties (Figure 1a). SHARP is further guided to perform actions optimal for molecular synthesis by action masks derived from pre-trained Synthesizability Estimation Models (SEM) (Figure 1b). Here, the action mask refers to the region specifications (e.g., bond, atom, or fragment) where an action should occur. Our action space is built in a two-level hierarchy: the first-level action selects the type of the action among diverse fragment-level movements such as substitution, addition, and deletion (Figure 1c). The second-level action selects which fragment to utilize for the designated action type. Fragments are chosen from a pre-generated dataset derived by BRICS decomposition onto actual chemical libraries to avoid the generation of invalid molecules (Figure 1d). By combining these hierarchical actions with an autoregressive generation pattern, SHARP supports the optimization of molecules through organic reaction pathways, providing a robust foundation for synthesizability,^16^ rather than relying on uncertain metrics like the SA score.^17^

**Figure 1:**
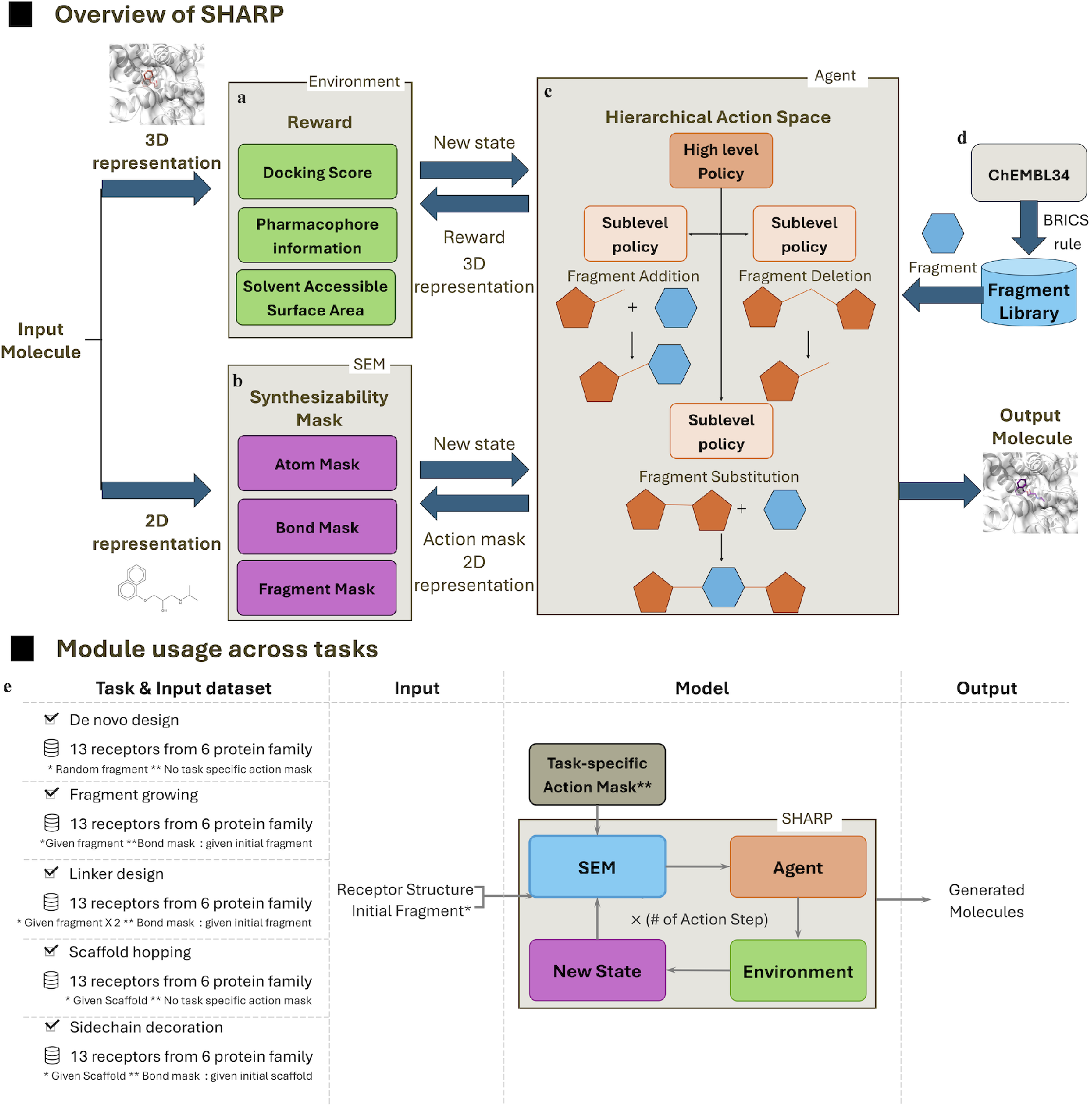
Overview of the SHARP. SHARP, a molecule generation model, was trained using a reinforcement learning framework. a) Policy networks were trained with rewards incorporating docking scores, pharmacophore information, and solvent-accessible surface area. b) Synthesizability masks were provided by the Synthesizability Estimation Model (SEM). c) Fragment-based hierarchical actions—addition, deletion, and substitution—were employed. d) A fragment library was created from ChEMBL34. e) Overview of SHARP module usage across tasks: De novo Design, Fragment Growing, Linker Design, Scaffold Hopping, Sidechain Decoration. All tasks were benchmarked using a high-binding-affinity molecule dataset from 13 receptors across six protein families.

The effectiveness of our approach, which a reinforcement learning agent drives, is rooted in three core components designed to discover molecules on the optimal Pareto front of synthesizability and binding affinity. The first is a versatile, hierarchical action space that allows not only molecular growth but also revisions during the generation process.^18^ Actions such as fragment addition and deletion can drive broad, global exploration, enabling the discovery of diverse and novel molecular scaffolds. On the other hand, fragment substitution facilitates fine-grained local optimization, allowing for minimal modifications to refine the properties of a promising candidate. This strategic division of actions ensures both compelling exploration of the vast chemical space and precise exploitation of promising regions. The second component is the reward function. It can effectively enhance the binding affinity of specific generated molecule by integrating docking scores and pharmacophore interactions. Our approach overcomes the respective challenges of reward sparsity in pharmacophore-only methods and the generation of giant molecules in docking score-only optimization;^19^ by synergistically integrating pharmacophore interactions, docking scores, and solvent-accessible surface area into a unified reward function, we enable a balanced optimization towards drug-like candidates with high predicted binding affinity. The third component is a dedicated Synthesizability Estimation Model (SEM), which addresses the other key objective by providing explicit feedback on the synthetic viability of each candidate. As our analyses demonstrate, the integration of these components within an RL framework proves highly effective at identifying novel, potent, and synthesizable molecules, successfully tackling the core challenges of global optimization in drug design.

One of the strengths of the SHARP method is that, it can be readily applied to a broad type of specific lead optimization tasks in addition to the original de novo design tasks (Figure 1 e) This adaptability is naturally realized by combining modules for customized goals and by defining specific action masks to constrain the generative process. Importantly, the validity of the generated molecules from various adaptations is guaranteed by the SEM module. In the following sections, we mainly report the performance based on the most generic de novo design scenario. We then present applications including de novo design, fragment growing, linker design, scaffold hopping, and sidechain decoration, with detailed case studies.

### Global optimization performance over multiple objectives

To provide practical guidance for benchmarking molecular generative models, we propose a new evaluation strategy using a newly curated test set and improved metrics. Our test set contains 13 diverse receptors with no redundancy, which contrasts with the CrossDock data set commonly used in previous related works. To evaluate the diverse perspectives of generated molecules, we included the Vina score, SC Score, and QED Score to measure potency, synthesizability, and drug-likeness, respectively. The most distinct evaluation metric introduced in this work is the “similarity” metric, which measures how closely the sampled molecules mimic one of the known binders to the receptor. We introduced this metric to evaluate the “real-world” success, as other metrics are too indirect to accurately represent the actual drug design success (for instance, molecules with a high Vina score do not guarantee tight binders). Details of the test set and evaluation metrics are provided in the Methods.

The performance of SHARP was benchmarked against several methods, including Pocket2Mol,^12^ TargetDiff,^15^ and DiffBP.^14^ As illustrated in Table 1, SHARP consistently generates molecules with the best Vina scores and high similarities to high-binding-affinity compounds, demonstrating superior performance with high statistical significance (*p <* 0.001). When measuring the proportion of molecules that improved upon a reference set (149 protein-ligand complex structures from PDBbind2020), SHARP outperformed all competitors, with 30.2% of its molecules showing a better Vina score and 2.7% showing higher similarity. SHARP exhibits comparable or superior distributions for both metrics with strong statistical significance (*p <* 0.005). Notably, 72.7% of SHARP’s molecules achieved an SC score below the 4.0 threshold for synthesizable compounds, a result competitive with one of the top-performing models in that category, DiffBP (80.5%).

**Table 1:** Quantitative performance comparison in molecular generation. This table evaluates the properties of molecules from various models, organized into key categories. Our model demonstrates statistically significant improvements (*p <* 0.001) in binding affinity (Vina Score, Similarity) and chemical viability (SC Score, QED) compared to other methods. Success and Recovery Rates measure superiority to reference molecules, while the Synthesizability Rate identifies molecules with an SC Score below 4.0, indicating ease of synthesis.

| Model Name | Binding affinity |  |  |  | Chemical Viability |  |  |
| --- | --- | --- | --- | --- | --- | --- | --- |
| | Vina Score (↓) | Success rate ($\%$, ↑) | Similarity (↑) | Recovery rate ($\%$, ↑) | SC score (↓) | Synthesizability rate ($\%$, ↑) | QED (↑) |
| SHARP | **-8.527 ± 1.522** | **30.2** | 0.205 ± 0.128 | **2.7** | 3.693 ± 0.467 | 72.7 | 0.610 ± 0.169 |
| Pocket2Mol | -6.437 ± 2.099 | 20.5 | 0.118 ± 0.044 | 0.0 | 3.697 ± 0.601 | 65.9 | 0.253 ± 0.157 |
| TargetDiff | -6.960 ± 2.198 | 12.8 | 0.143 ± 0.037 | 0.0 | 3.876 ± 0.659 | 53.6 | 0.506 ± 0.208 |
| GraphBP | -5.374 ± 1.329 | 2.9 | 0.106 ± 0.038 | 0.0 | **2.922 ± 0.493** | **97.3** | 0.491 ± 0.150 |
| DiffSBDD | -5.924 ± 2.298 | 7.2 | 0.121 ± 0.031 | 0.0 | 3.812 ± 0.627 | 58.1 | 0.323 ± 0.188 |
| LIGAN | -7.640 ± 1.647 | 14.2 | **0.381 ± 0.282** | 0.3 | 3.488 ± 0.539 | 75.5 | **0.679 ± 0.197** |
| DiffBP | -6.237 ± 1.739 | 8.5 | 0.112 ± 0.033 | 0.0 | 3.561 ± 0.532 | 80.5 | 0.356 ± 0.166 |

We then performed a Pareto front analysis to validate the optimization performances across methods thoroughly. We mainly investigated the trade-off between potency and synthesizability (Figure 2). The analysis, which plots Vina score and similarity against SC score, confirms that SHARP’s Pareto front consistently surpasses those of all other methods. This result demonstrates that SHARP achieves superior potency without compromising synthetic feasibility, finding a more optimal balance between these conflicting objectives.

**Figure 2:**
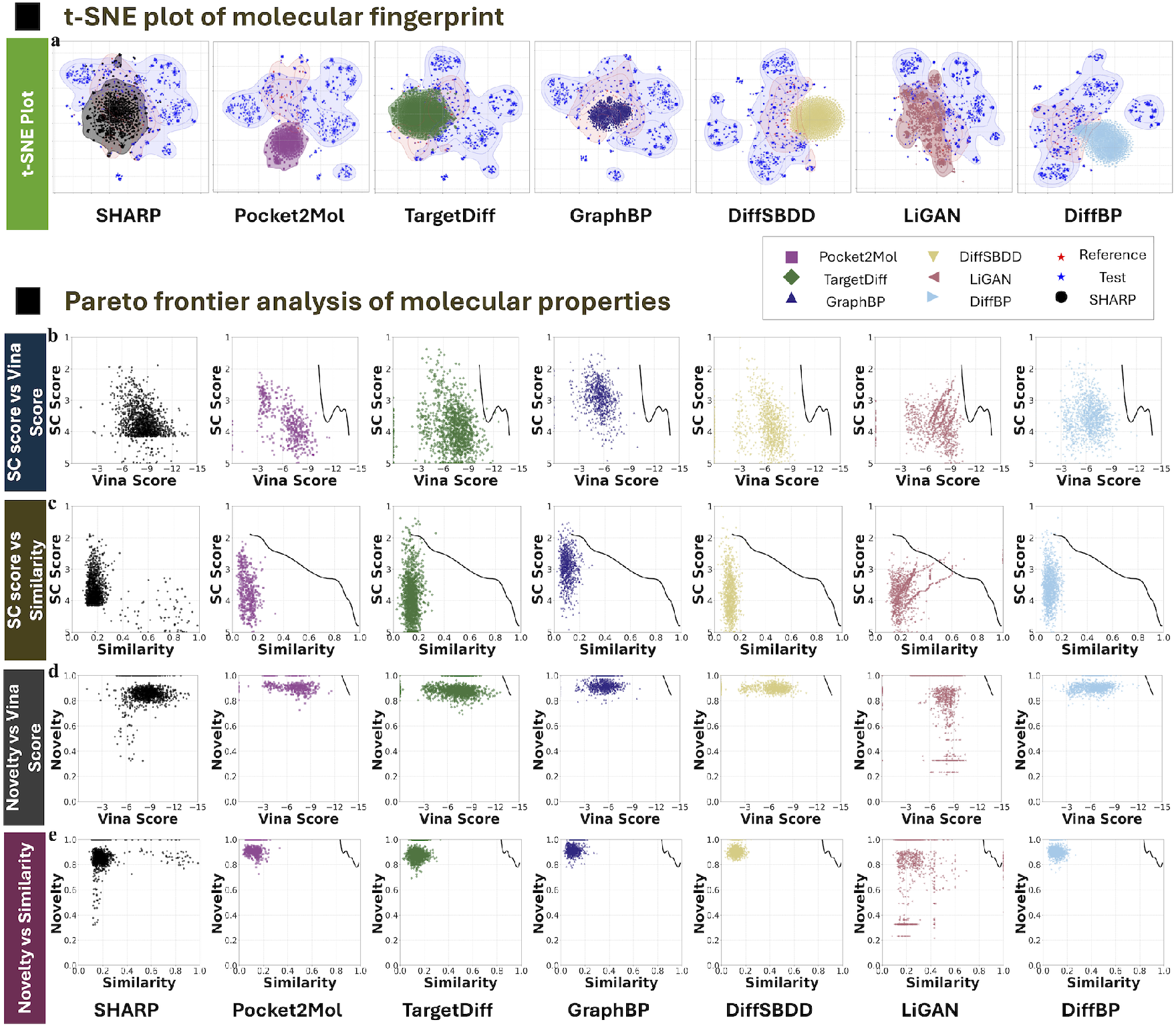
t-SNE and Pareto frontier analysis. (a) t-SNE Plots of Molecular Distributions: These t-SNE plots visualize the molecular distributions from each model, as well as the training and test sets. The SHARP model shows a notably sparse distribution, distinct from the reference set yet closer to the test set, indicating generation of high-binding affinity, structurally novel molecules. To verify that our model (SHARP) optimizes potency without sacrificing other key metrics, Pareto fronts are plotted for: (b) SC score vs. Vina Score, (c) SC score vs. Similarity, (d) Novelty vs. Vina Score, and (e) Novelty vs. Similarity, for each evaluated model. For comparative ease, SHARP’s Pareto front is additionally depicted as a thick black line on each other model’s plot. The Vina score axis is inverted such that compounds exhibiting both high novelty and high potency are positioned in the top-right quadrant. For clarity, SHARP’s Pareto front is overlaid as a thick black line on the plots of other models. Consistently, SHARP’s Pareto front is positioned superiorly across all comparisons, demonstrating that our model achieves potency optimization without compromising SC score or novelty. This superior performance is attributed to the introduced fragment-based optimization and SEM.

We hypothesize that these notable results stem from the synergy of SHARP’s core components. The high potency is attributed to the combination of broad chemical space exploration, enabled by fragment-based generation, and efficient exploitation, facilitated by hierarchical actions. This is further enhanced by a reward function that incorporates pharmacophore interaction and SASA information, resulting in more accurate predictions of real-world potency compared to relying solely on Vina scores. Concurrently, strong synthesizability and drug-likeness are maintained by the use of a rational fragment library and the implicit guidance of the Synthesizability Estimation Model (SEM). We claim that implicit treatment of synthesizability via SEM was more effective than introducing an explicit reward term; conceptually, this allows for avoiding over-complicating the search landscape and hence can effectively lead the optimization process to Pareto optima.

### Scaffold exploration and structural diversity beyond reference data

The internal diversity and novelty of generated molecules by SHARP are assessed and reported in Table 2. The diversity was evaluated at both the 2D intramolecular fingerprint level and the 3D protein-ligand interaction level (details in Methods). For 2D fingerprint diversity, as shown in Table 2, the model demonstrates statistically significant (*p <* 0.001) comparable or higher diversity in Tanimoto distance and achieved a highly competitive 88.1% ratio of unique Bemis-Murcko scaffolds, ^20^ close to TargetDiff’s leading 90.0%. Similarly, for 3D interaction diversity, SHARP exhibits an excellent distribution of protein-ligand interaction fingerprints (*p <* 0.001).

**Table 2:** Diversity and novelty comparison in molecular generation. This table evaluates molecular diversity and novelty using two methodologies: 2D chemical fingerprints and 3D interaction fingerprints. First, using 2D fingerprints, we assess Diversity (based on internal fingerprint distances and the percentage of unique scaffolds) and Novelty (based on distances to the reference set and the percentage of new scaffolds). In these metrics, our model demonstrates superior performance. This analysis is then repeated using 3D interaction fingerprints, where our model achieves comparable or superior results again. Together, these findings confirm our model’s strong capacity for extensively exploring both the chemical and interaction space.

| Model name | Fingerprint Diversity (↑) | Fingerprint Novelty (↑) | Scaffold Diversity(↑) | Scaffold Novelty(↑) | Interaction Diversity(↑) | Interaction Novelty (↑) |
| --- | --- | --- | --- | --- | --- | --- |
| SHARP | 0.874 ± 0.046 | 0.867 ± 0.084 | 88.1 | **98.7** | 0.744 ± 0.190 | 0.976 ± 0.087 |
| Pocket2Mol | 0.878 ± 0.056 | 0.921 ± 0.047 | 55.9 | 76.8 | **0.809 ± 0.142** | 0.919 ± 0.191 |
| TargetDiff | **0.908 ± 0.031** | 0.894 ± 0.055 | **90.0** | 94.7 | 0.761 ± 0.188 | **0.979 ± 0.078** |
| GraphBP | 0.893 ± 0.072 | **0.928 ± 0.042** | 29.5 | 55.2 | 0.764 ± 0.187 | 0.968 ± 0.119 |
| DiffSBDD | 0.909 ± 0.035 | 0.914 ± 0.046 | 65.8 | 78.2 | 0.774 ± 0.184 | 0.976 ± 0.084 |
| LiGAN | 0.850 ± 0.179 | 0.792 ± 0.212 | 54.5 | 97.1 | 0.727 ± 0.229 | 0.914 ± 0.238 |
| DiffBP | 0.879 ± 0.051 | 0.916 ± 0.044 | 49.0 | 79.4 | 0.741 ± 0.200 | 0.963 ± 0.130 |

The novelty of molecules generated by SHARP was confirmed through 2D, 3D, and visual chemical space analyses. For the 2D fingerprint novelty, SHARP demonstrates remarkably high novelty with statistical significance (*p <* 0.001) and achieves the highest value of 98.7% for generating Bemis-Murcko scaffolds not present in the reference set. This high novelty was also observed in 3D protein-ligand interactions, where the model again demonstrated excellent novelty (*p <* 0.001). The definition for these “unique” interactions is provided in Section. Finally, these findings are visually confirmed by t-SNE plots in Figure 2a, which show SHARP exploring a significantly larger chemical space, closer to the test set and more distant from the reference data than other methods. We hypothesize that this superior diversity and novelty originate from our fragment-based generation, which facilitates extensive exploration of chemical space.

Pareto analyses on the novelty with other metrics are shown in Figure 2c and Figure 2d. As shown in the figures, SHARP’s novelty is not a simple outcome of a trade-off with other metrics, but rather emerges as other metrics are improved simultaneously. Pareto front of the molecules generated by SHARP consistently surpasses those of other methods, demonstrating its multi-objective performance.

### In-depth Analysis of SHARP: Optimization Process and Component Contributions

In this section, we analyze the origins of SHARP’s performance. We first analyze the model’s behavior during the molecular generation process, followed by the evaluation of the importance of core modules through an ablation study. Detailed methodologies for these analyses are presented in method sections.

Our investigation began with a detailed analysis of the Reinforcement Learning (RL) agent’s behavior. As illustrated in Figure 3a, the docking reward shows a statistically significant increase (Pearson correlation coefficient of 0.392, *p <* 0.001) as generation steps advance, validating that SHARP effectively optimizes molecules over time. To ensure this was not due to overfitting, we analyzed fragment usage patterns (Figure 3b). The model utilizes a more diverse set of fragments than the training data, as confirmed by a higher Shannon entropy (4.646 vs. 2.754) and a greater Jaccard similarity to the test set, implying that the model has learned actions beyond the training set.(*p <* 0.001) An analysis of action utilization (Figure 3c) shows that fragment substitutions are dominant in our molecule generation. The nature of this action evolves, with the model making size-matched substitutions in later steps (Figure 3d), which is consistent with an exploitation phase of optimization. An example trajectory (Figure 3e) further demonstrates the agent’s capacity for global optimization, as it accepts a decrease in short-term reward to achieve long-term reward, which leads to superior final molecules.

**Figure 3:**
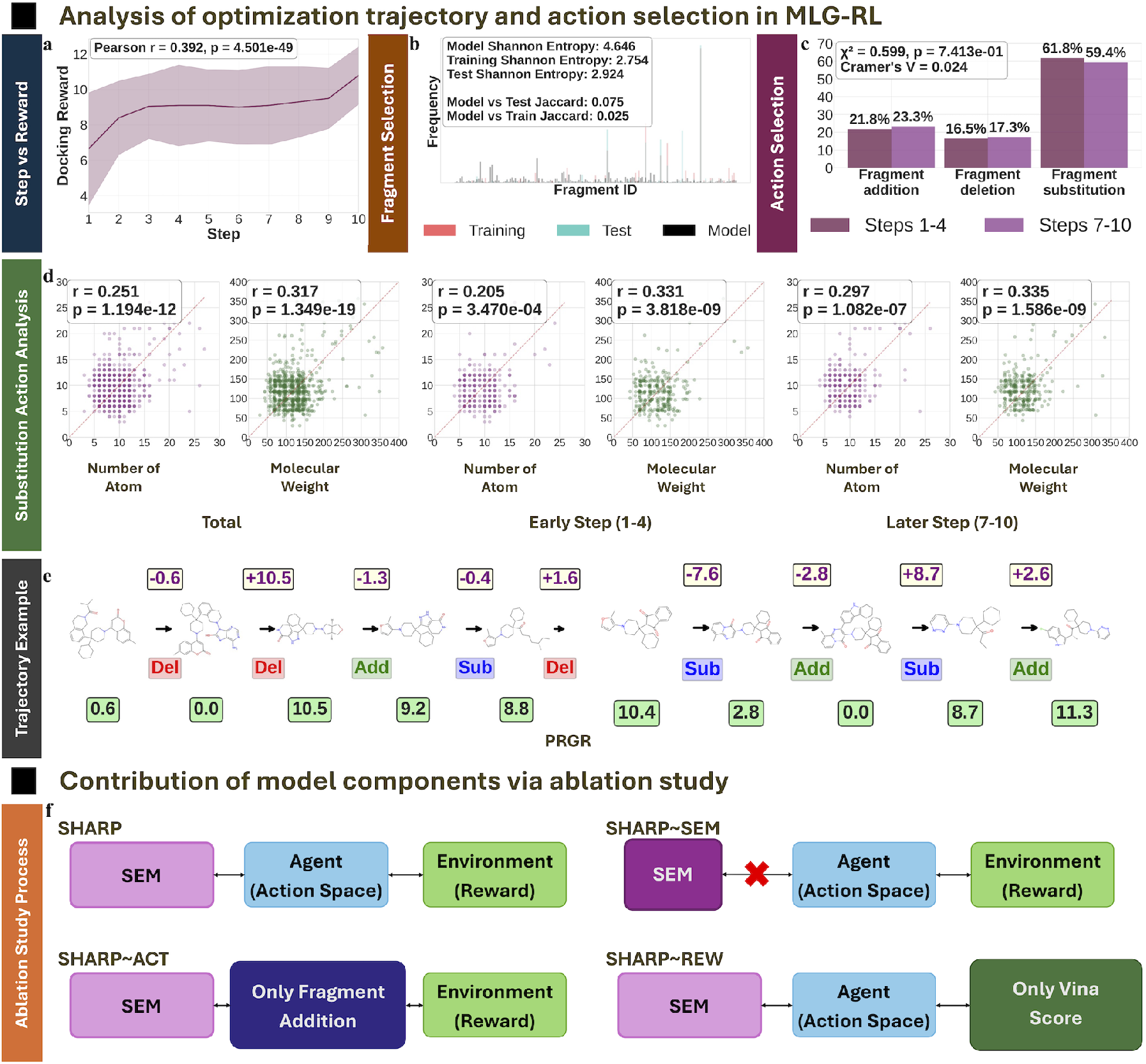
Analyzing action selection and contribution of each modules in SHARP. (a) Illustrates the steady increase in rewards during generation. The solid line and shaded region represent the median and standard deviation, respectively. (b) This metric displays the model’s diverse fragment selection, which differs from the training data, indicating that there is no overfitting. For clarity, only fragments present in all sets are displayed. (c) Compares action distribution between early (1-4) and late (7-10) generation steps, revealing a consistent utilization pattern. (d) Shows that in later steps, the model uses similarly-sized fragments for substitution, suggesting a strategy of making minor modifications to exploit the chemical space. (e) An example for the PRGR protein, demonstrating the model’s ability to sacrifice short-term rewards for global optimization to achieve a better final molecule. (f) Describes the complete SHARP model architecture alongside its ablative versions, which lack specific modules.

Next, to assess the contribution of each model component, we conducted an ablation study with three variants as described in Figure 3f: SHARP~SEM (lacking the Synthesizability Estimation Model), SHARP~ACT (restricted to fragment addition actions), and SHARP~REW (using only the Vina docking score for its reward). As shown in Table 3, these variants exhibited distinct trade-offs. SHARP~SEM achieved the best Vina Score but lower similarity, while the addition-only SHARP~ACT excelled in synthesizability (SC score) and drug-likeness (QED score) at the cost of effective optimization. The simplified-reward SHARP~REW model performed poorly across similarity, synthesizability, and drug-likeness. These results provide several key insights about the contributions of modules. First, the performance of SHARP~SEM suggests the SEM module acts as a helpful constraint, guiding optimization towards synthesizable molecules, albeit with a minor trade-off in similarity. Second, the SHARP~REW variant, which used a Vina-score-only reward, produced molecules with poorer similarity, synthesizability, and drug-likeness, confirming that this simplified reward function tends to generate larger, less drug-like compounds. Third, the poor global optimization of SHARP~ACT highlights that fragment deletion and substitution actions are crucial for effective optimization, enabling fine-tuning and correcting initial fragment placements. Finally, the consistent diversity and novelty across all variants highlight that our core fragment-based generation mechanism is the primary driver for exploring a vast chemical space.

**Table 3:**
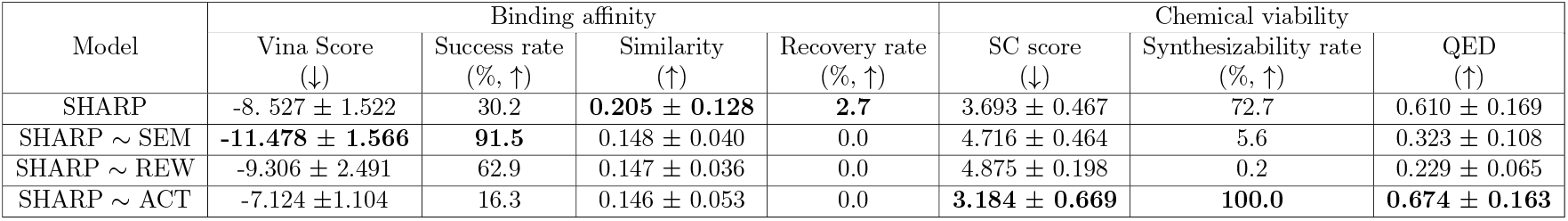
Performance comparison of SHARP and various versions of SHARP without specific module. The evaluation metrics are grouped into three categories: binding affinity (Vina Score, Success Rate, Similarity, Recovery Rate), synthesizability (SC Score, Synthesizability Rate), and drug-likeness (QED). Detailed definitions for each metric are provided in the Methods section.

| Model | Binding affinity |  |  |  | Chemical viability |  |  |
| --- | --- | --- | --- | --- | --- | --- | --- |
|  | Vina Score<br>(↓) | Success rate<br>(%, ↑) | Similarity<br>(↑) | Recovery rate<br>(%, ↑) | SC score<br>(↓) | Synthesizability rate<br>(%, ↑) | QED<br>(↑) |
| SHARP | -8.527 ± 1.522 | 30.2 | **0.205 ± 0.128** | **2.7** | 3.693 ± 0.467 | 72.7 | 0.610 ± 0.169 |
| SHARP ~ SEM | **-11.478 ± 1.566** | **91.5** | 0.148 ± 0.040 | 0.0 | 4.716 ± 0.464 | 5.6 | 0.323 ± 0.108 |
| SHARP ~ REW | -9.306 ± 2.491 | 62.9 | 0.147 ± 0.036 | 0.0 | 4.875 ± 0.198 | 0.2 | 0.229 ± 0.065 |
| SHARP ~ ACT | -7.124 ± 1.104 | 16.3 | 0.146 ± 0.053 | 0.0 | **3.184 ± 0.669** | **100.0** | **0.674 ± 0.163** |

### A Retrospective Validation of SHARP: Analysis of Molecular Rationality

We investigated whether molecules generated by SHARP contain strained or lengthy macro-cycles, which are observed in real drugs but can negatively impact stability and synthesizability. As shown in Figure 4a-d, SHARP consistently produced the lowest percentage of molecules containing 3-membered, 4-membered, 7-membered, and macrocyclic rings compared to other methods. This indicates a significantly reduced propensity to generate substructures that are hard to synthesize or less stable, a success that we attribute to our fragment library. To provide a reference, we analyzed the percentages of molecules containing 3-membered, 4-membered, 7-membered, and macrocyclic rings in the ChEMBL 34 dataset. The results show low percentages for each of these ring categories, which supports the assertion that such ring sizes are uncommon. This finding further substantiates that the SHARP model generates realistic, drug-like molecules.

**Figure 4:**
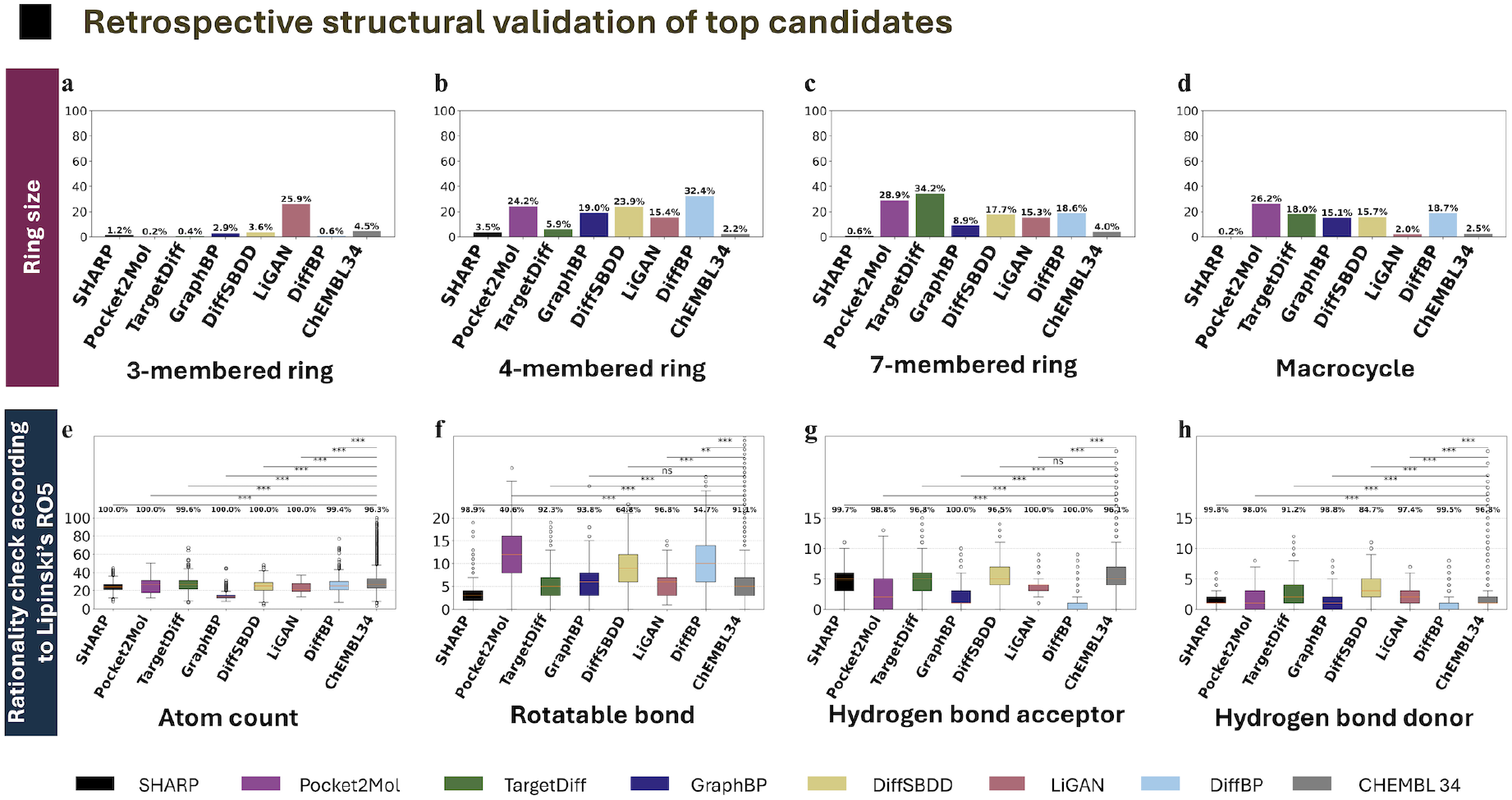
Rationality evaluation of generated molecules. Molecules from various methods, including SHARP (color-coded by method type in the lower panel), are analyzed for substructure, showing the percentage of molecules containing (a) 3-membered rings, (b) 4-membered rings, (c) 7-membered rings, and (d) macrocycles (rings with more than seven members). Additional generated molecule statistics include (e) atom count, (f) number of rotatable bonds, (g) hydrogen bond acceptors, and (h) hydrogen bond donors. Results are compared with ChEMBL34 statistics. We note that the asterisks indicate the level of statistical significance in the results (ns for *p >* 0.05, * for *p <* 0.05, ** for *p <* 0.01, and *** for *p <* 0.001).

We then examined key physicochemical properties, using the ChEMBL34 database^21^ as a reference for drug-like molecules. Excessive atom counts, rotatable bonds, or hydrogen bond donors/acceptors can compromise binding affinity and bioavailability. Distributions for atom count and rotatable bonds are highly comparable to those of ChEMBL34, avoiding the common pitfalls of other methods that generate molecules that are either too small or overly flexible (Figures 4e-f). Similarly, the distributions for hydrogen bond acceptors and donors align well with the reference and remain within the acceptable limits defined by Lipinski’s Rule of Five^22^ (Figures 4g-h). Consequently, SHARP demonstrates Lipinski compliance values that are either the highest or comparable to the top-performing methods. We hypothesize this is due to both the valid fragment library and the inclusion of pharmacophore and Solvent Accessible Surface Area (SASA) information in the reward function, which prevents the generation of excessively large molecules.

### Generalizability across multiple lead optimization tasks

To benchmark its general applicability, we evaluated SHARP’s performance against established methods (e.g., Pocket2Mol, TargetDiff) across four distinct lead optimization scenarios: fragment growing, linker design, scaffold hopping, and sidechain decoration. To describe details, fragment growing addresses scenarios where a promising fragment with key pharmacophores interacts with specific binding pocket residues. Linker design involves connecting multiple fragments exhibiting key pharmacophore interactions in different binding pocket regions. Scaffold hopping focuses on identifying and substituting the central core of existing active molecules.^23,24^ Sidechain decoration, the final task, aims to enhance binding affinity. For each task, SHARP employed a task-specific action mask combined with its Synthesizability Estimation Model (SEM). This approach guided the generative agent by preventing modification of predefined starting points (e.g., initial fragments or Bemis-Murcko scaffolds), ensuring task-relevant and synthesizable molecular design.

Across all four tasks, SHARP consistently generated molecules with high binding affinity and superior mean similarity to known active compounds (Table 4). Its performance was particularly outstanding in scaffold hopping, where it achieved a Vina score success rate(better Vina score than reference) of 40.6% and a recovery rate(Tanimoto similarity *>* 0.6 with high binding affinity molecules) of 42.8%, significantly outperforming TargetDiff (33.6% and 14.4%, respectively). In sidechain decoration, SHARP remained highly competitive with a 46.7% Vina score percentage and 50.5% similarity. Furthermore, molecules generated by SHARP consistently exhibited reasonable synthesizability (SC scores *<* 4.0) and drug-like properties (QED), showing notable synthesizability in fragment growing and linker design compared to other high-performing models.

**Table 4:**
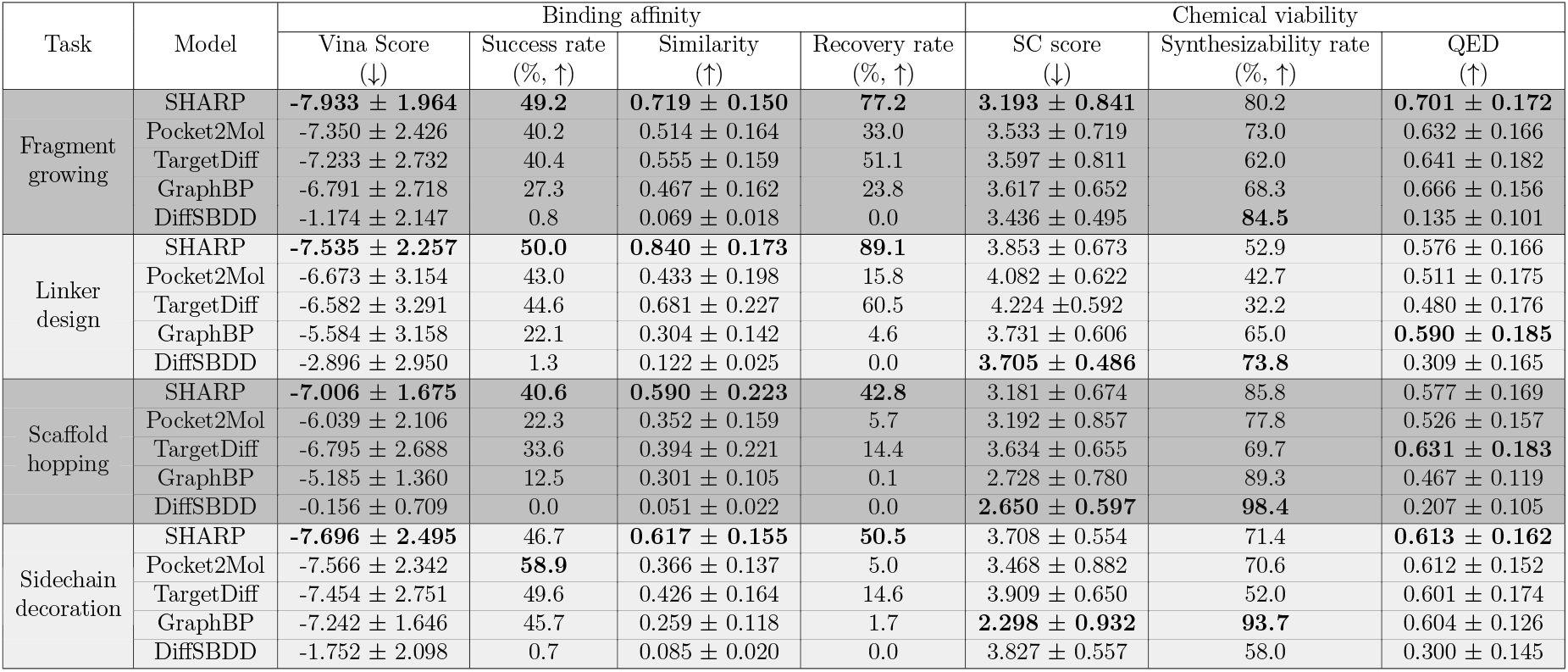
Quantitative performance comparison in various lead optimization tasks. Performance comparison of SHARP against other generative models across four lead optimization tasks: fragment growing, linker design, scaffold hopping, and sidechain decoration. Models were evaluated for their optimization ability using the Vina Score and Similarity, and for chemical viability using the SC Score and QED. The reported rates indicate the percentage of molecules exceeding a threshold: Success Rate (better Vina score than reference), Recovery Rate (Tanimoto similarity *>* 0.6 with high binding affinity molecules), and Synthesizability Rate (SC score *<* 4.0). SHARP demonstrates comparable or superior performance across all tasks and metrics.

| Task | Model | Binding affinity |  |  |  | Chemical viability |  |  |
| --- | --- | --- | --- | --- | --- | --- | --- | --- |
|  |  | Vina Score<br>(↓) | Success rate<br>(%, ↑) | Similarity<br>(↑) | Recovery rate<br>(%, ↑) | SC score<br>(↓) | Synthesizability rate<br>(%, ↑) | QED<br>(↑) |
| Fragment growing | SHARP | **-7.933 ± 1.964** | **49.2** | **0.719 ± 0.150** | **77.2** | **3.193 ± 0.841** | 80.2 | **0.701 ± 0.172** |
|  | Pocket2Mol | -7.350 ± 2.426 | 40.2 | 0.514 ± 0.164 | 33.0 | 3.533 ± 0.719 | 73.0 | 0.632 ± 0.166 |
|  | TargetDiff | -7.233 ± 2.732 | 40.4 | 0.555 ± 0.159 | 51.1 | 3.597 ± 0.811 | 62.0 | 0.641 ± 0.182 |
|  | GraphBP | -6.791 ± 2.718 | 27.3 | 0.467 ± 0.162 | 23.8 | 3.617 ± 0.652 | 68.3 | 0.666 ± 0.156 |
|  | DiffSBDD | -1.174 ± 2.147 | 0.8 | 0.069 ± 0.018 | 0.0 | 3.436 ± 0.495 | **84.5** | 0.135 ± 0.101 |
| Linker design | SHARP | **-7.535 ± 2.257** | **50.0** | **0.840 ± 0.173** | **89.1** | 3.853 ± 0.673 | 52.9 | 0.576 ± 0.166 |
|  | Pocket2Mol | -6.673 ± 3.154 | 43.0 | 0.433 ± 0.198 | 15.8 | 4.082 ± 0.622 | 42.7 | 0.511 ± 0.175 |
|  | TargetDiff | -6.582 ± 3.291 | 44.6 | 0.681 ± 0.227 | 60.5 | 4.224 ± 0.592 | 32.2 | 0.480 ± 0.176 |
|  | GraphBP | -5.584 ± 3.158 | 22.1 | 0.304 ± 0.142 | 4.6 | 3.731 ± 0.606 | 65.0 | **0.590 ± 0.185** |
|  | DiffSBDD | -2.896 ± 2.950 | 1.3 | 0.122 ± 0.025 | 0.0 | **3.705 ± 0.486** | **73.8** | 0.309 ± 0.165 |
| Scaffold hopping | SHARP | **-7.006 ± 1.675** | **40.6** | **0.590 ± 0.223** | **42.8** | 3.181 ± 0.674 | 85.8 | 0.577 ± 0.169 |
|  | Pocket2Mol | -6.039 ± 2.106 | 22.3 | 0.352 ± 0.159 | 5.7 | 3.192 ± 0.857 | 77.8 | 0.526 ± 0.157 |
|  | TargetDiff | -6.795 ± 2.688 | 33.6 | 0.394 ± 0.221 | 14.4 | 3.634 ± 0.655 | 69.7 | **0.631 ± 0.183** |
|  | GraphBP | -5.185 ± 1.360 | 12.5 | 0.301 ± 0.105 | 0.1 | 2.728 ± 0.780 | 89.3 | 0.467 ± 0.119 |
|  | DiffSBDD | -0.156 ± 0.709 | 0.0 | 0.051 ± 0.022 | 0.0 | **2.650 ± 0.597** | **98.4** | 0.207 ± 0.105 |
| Sidechain decoration | SHARP | **-7.696 ± 2.495** | 46.7 | **0.617 ± 0.155** | **50.5** | 3.708 ± 0.554 | 71.4 | **0.613 ± 0.162** |
|  | Pocket2Mol | -7.566 ± 2.342 | **58.9** | 0.366 ± 0.137 | 5.0 | 3.468 ± 0.882 | 70.6 | 0.612 ± 0.152 |
|  | TargetDiff | -7.454 ± 2.751 | 49.6 | 0.426 ± 0.164 | 14.6 | 3.909 ± 0.650 | 52.0 | 0.601 ± 0.174 |
|  | GraphBP | -7.242 ± 1.646 | 45.7 | 0.259 ± 0.118 | 1.7 | **2.298 ± 0.932** | **93.7** | 0.604 ± 0.126 |
|  | DiffSBDD | -1.752 ± 2.098 | 0.7 | 0.085 ± 0.020 | 0.0 | 3.827 ± 0.557 | 58.0 | 0.300 ± 0.145 |

SHARP’s high generalizability across these diverse tasks is directly attributable to the synergistic combination of its core components. The hierarchical action space, with its unique fragment deletion and substitution capabilities, was critical for its superior performance in complex tasks like scaffold hopping. This same action space enables SHARP to effectively perform fine-grained modifications for sidechain decoration, a domain typically dominated by atom-based models. Finally, the integrated SEM ensures this generative flexibility does not compromise synthesizability, allowing SHARP to effectively build upon promising starting fragments while generating viable molecules.

### A Retrospective Validation of SHARP: Case study

In the following section, we demonstrate the specific application of SHARP for solving practical problems on four distinct lead optimization tasks.

In the PRTM5 MTA complex system, ^25^ the goal was to grow a starting fragment to maximize hydrophobic interactions within the binding site. As illustrated in Figure 5a, SHARP’s top molecules successfully fit the binding site’s shape while also establishing a key polar interaction with GLN 309, demonstrating a nuanced understanding of the local chemical environment.

**Figure 5:**
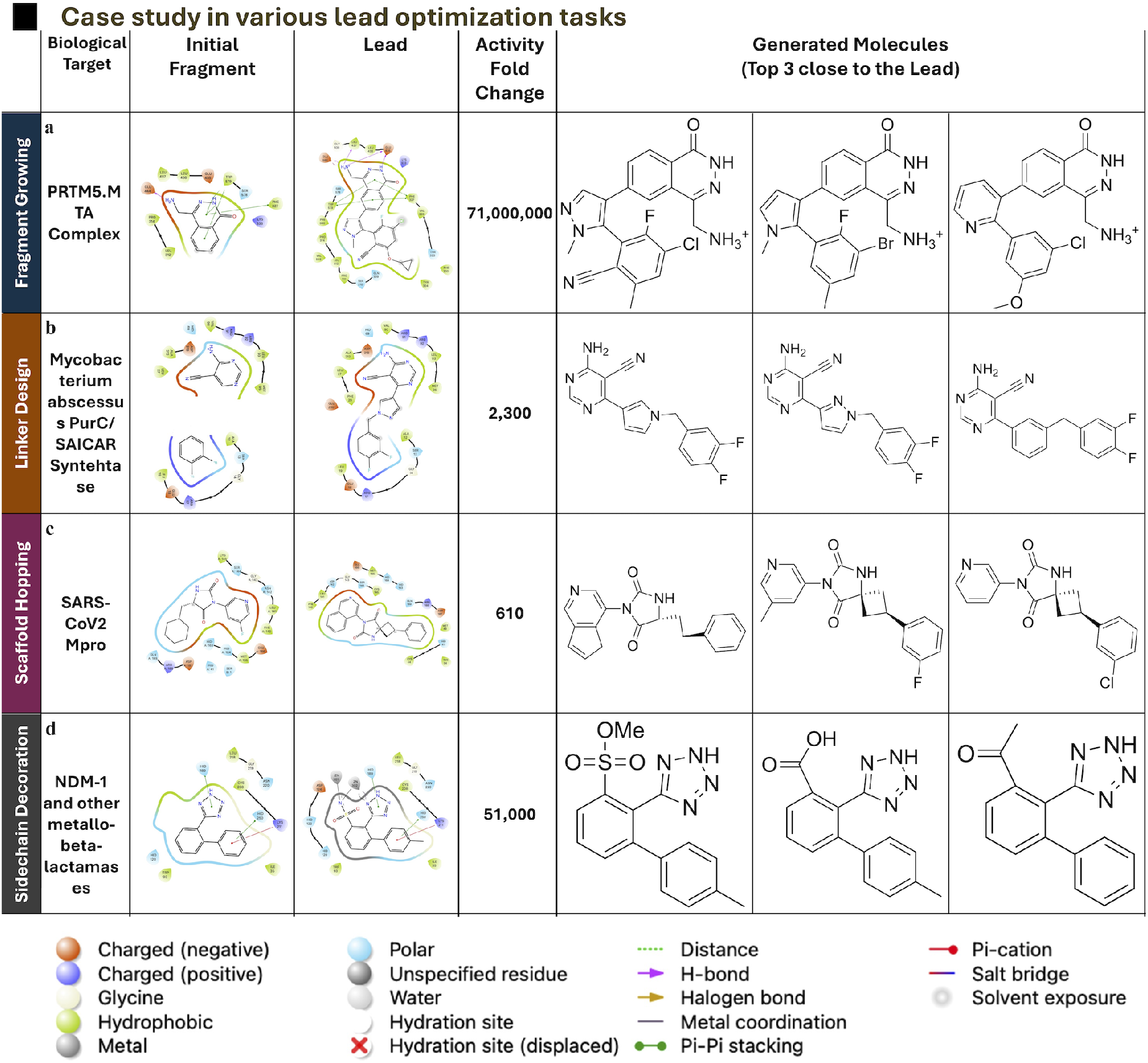
Case study in various lead optimization tasks. This figure presents a case study for four lead optimization tasks: (a) Fragment Growing, (b) Linker Design, (c) Scaffold Hopping, and (d) Sidechain Decoration. For each task, we selected a distinct biological system. The figure highlights the biological target, the protein-ligand interactions of the initial fragment, the interactions of the known lead compound, and the activity fold change. Crucially, we also display the top three molecules generated by SHARP that are most similar to the lead compound for each task. These results confirm SHARP’s ability to generate compounds structurally similar to known lead compounds across various lead optimization scenarios, even with different test sets.

For the Mycobacterium abscessus PurC/SAICAR Synthetase system,^26^ the objective was to design a rigid linker to minimize entropic penalties. SHARP’s top compounds incorporated ring substructures into the linker, successfully reducing rotatable bonds while maintaining an optimal orientation for key interactions (Figure 5b).

Using the SARS-CoV2 Mpro system,^27^ this task required modifying both ends of a molecule to improve receptor fit. The top compounds generated by SHARP exhibit an increased linker radius and larger fragments, enhancing complementarity with the binding site and confirming the model’s ability to solve complex geometric challenges (Figure 5c).

In the NDM-1 metallo-β-lactamase system,^28^ the crucial task was to introduce sidechains capable of forming polar interactions near a metal center. SHARP successfully identified and incorporated these necessary interactions in its top generated compounds, showcasing its accurate perception of the chemical environment (Figure 5d).

In summary, our retrospective validation confirms the robustness of the SHARP model. The molecular rationality analysis demonstrates that SHARP consistently generates structurally and physicochemically sound molecules satisfying well-known drug-likeness properties. Furthermore, case studies confirm that SHARP can successfully translate this capability into practical performance, characterizing binding site geometries and chemical environments to tackle a diverse range of lead optimization challenges. Together, these findings validate SHARP as an effective tool for structure-based drug design.

## Discussion

We introduced SHARP, a molecular generative model designed for generating and modifying high-binding-affinity molecules within the context of given receptor structures. SHARP frames the molecular generation task as a global optimization problem rather than solely a distribution modeling task. As validated through quantitative and generalizability analyses, its superior performance across de novo design and diverse lead optimization scenarios stems from the synergy of four critical components. These include a refined reward function that guides the generation of potent, real-world molecules and enables goal-oriented design; a Synthesizability Estimation Model (SEM) ensuring rational and synthesizable molecule generation, as proven by quantitative and rationality analyses; a fragment-based hierarchical action space providing a diverse set of operations for broad exploration in early stages and fine-tuned exploitation in later steps, evidenced by quantitative and action analyses, confirmed across all analyses. Our ablation study further demonstrated that all four modules are essential for achieving SHARP’s high performance. Addressing the well-documented limitations of prior fragment-based reinforcement learning models—such as the generation of excessively large, non-drug-like molecules from optimizing docking scores alone^29^, inefficient optimization due to the sparse nature of pharmacophoric rewards^30^, and poor exploitation from relying solely on fragment addition^31^ —our model employs a comprehensive strategy by combining a diverse action space (addition, deletion, and substitution) with a synergistic, multi-objective reward function to enable the balanced and robust generation of drug-like candidates.

SHARP’s capabilities could be further improved. We suggest enhancing two key modules: the reward function and the Synthesizability Estimation Model. Replacing the current reward function with a more accurate binding affinity estimation model would increase the likelihood of identifying molecules with true high binding affinity. Additionally, the current Synthesizability Estimation Model, which is based on BRICS rules, could be improved by training it on a broader range of organic reactions, making SHARP’s actions more representative of actual chemical synthesis. These enhancements, along with real-world applications to make a lead compound against specific targets, will be our work in the immediate future.

## Experimental Details

### Dataset

For training a reinforcement learning agent, we required receptor structures, which were obtained from the CrossDocked2020 dataset.^32^ We filtered the receptors based on two criteria: (i) binding site structural similarity with the test set exceeding a defined threshold and (ii) ligands classified as peptides. To evaluate binding site structural similarity, we utilized FLAPP,^33^ which aligns pockets from the training set with those in the test set and quantifies similarity using the ratio of successfully aligned amino acids. A threshold score of 0.6 was applied. Following this filtering process, we compiled a final list of 5,817 receptors. We split them into 8:2 for training and validation, which also ensures there is no similarity between the training and validation sets in terms of binding site structural similarity.

For our test set, we selected receptors from the DUD-E dataset,^34^ known for its well-classified protein families. The specific receptors chosen are detailed in Table 5. Notably, FKBP1A and Neprilysin are included, systems also present in the CrossDocked2020 test set, which distinguishes training and test sets based on sequence similarity. High-affinity binding molecules for each receptor were sourced from BindingDB,^35^ a comprehensive compilation of binding affinity assay data. The exact number of high-affinity molecules for each receptor is also provided in Table 5. To further prevent ligand information leakage, we filtered out test set molecules that shared a 2D fingerprint Tanimoto similarity exceeding 0.6 with any molecule in the reference set. The reference set consists fo the native ligands experimentally resolved within the three-dimensional structures of target protein-ligand complexes.

**Table 5:**
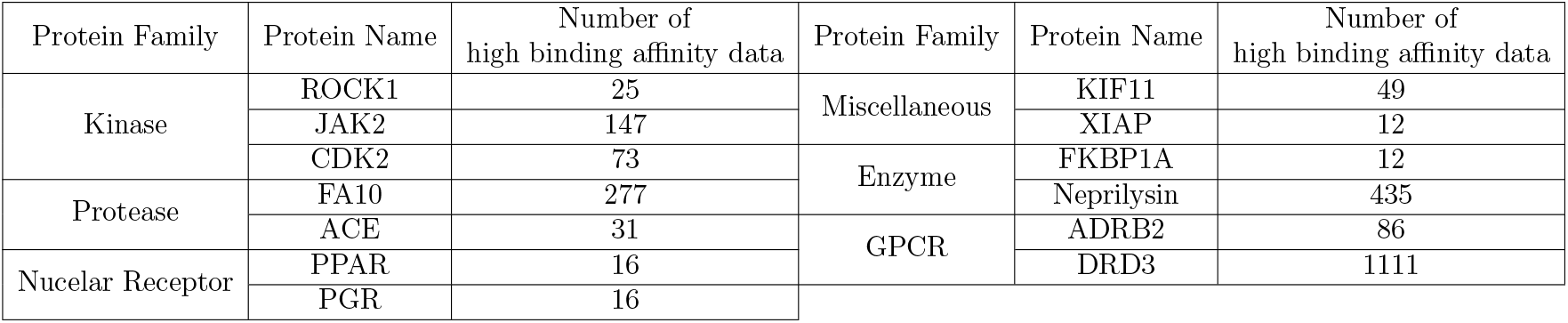
Selected protein targets across diverse families, with corresponding counts of high-affinity ligands.

| Protein Family | Protein Name | Number of high binding affinity data | Protein Family | Protein Name | Number of high binding affinity data |
| --- | --- | --- | --- | --- | --- |
| Kinase | ROCK1 | 25 | Miscellaneous | KIF11 | 49 |
|  | JAK2 | 147 |  | XIAP | 12 |
|  | CDK2 | 73 | Enzyme | FKBP1A | 12 |
| Protease | FA10 | 277 |  | Neprilysin | 435 |
|  | ACE | 31 | GPCR | ADRB2 | 86 |
| Nuclear Receptor | PPAR | 16 |  | DRD3 | 1111 |
|  | PGR | 16 |  |  |  |

We employed the BRICS fragmentation rule^36^ to construct the fragment library. Using this rule, we fragmented molecules from the ChEMBL34 dataset,^21^ and the resulting fragmentation logs served as the training set for the synthesizability estimation model.

### Model Architecture

SHARP is composed of two main components: the Synthesizability Estimation Model (SEM) and the Reinforcement Learning (RL) agent (Figure 6 and 7). Below, we first describe the functionality of the Synthesizability Estimation Model (SEM).

**Figure 6:**
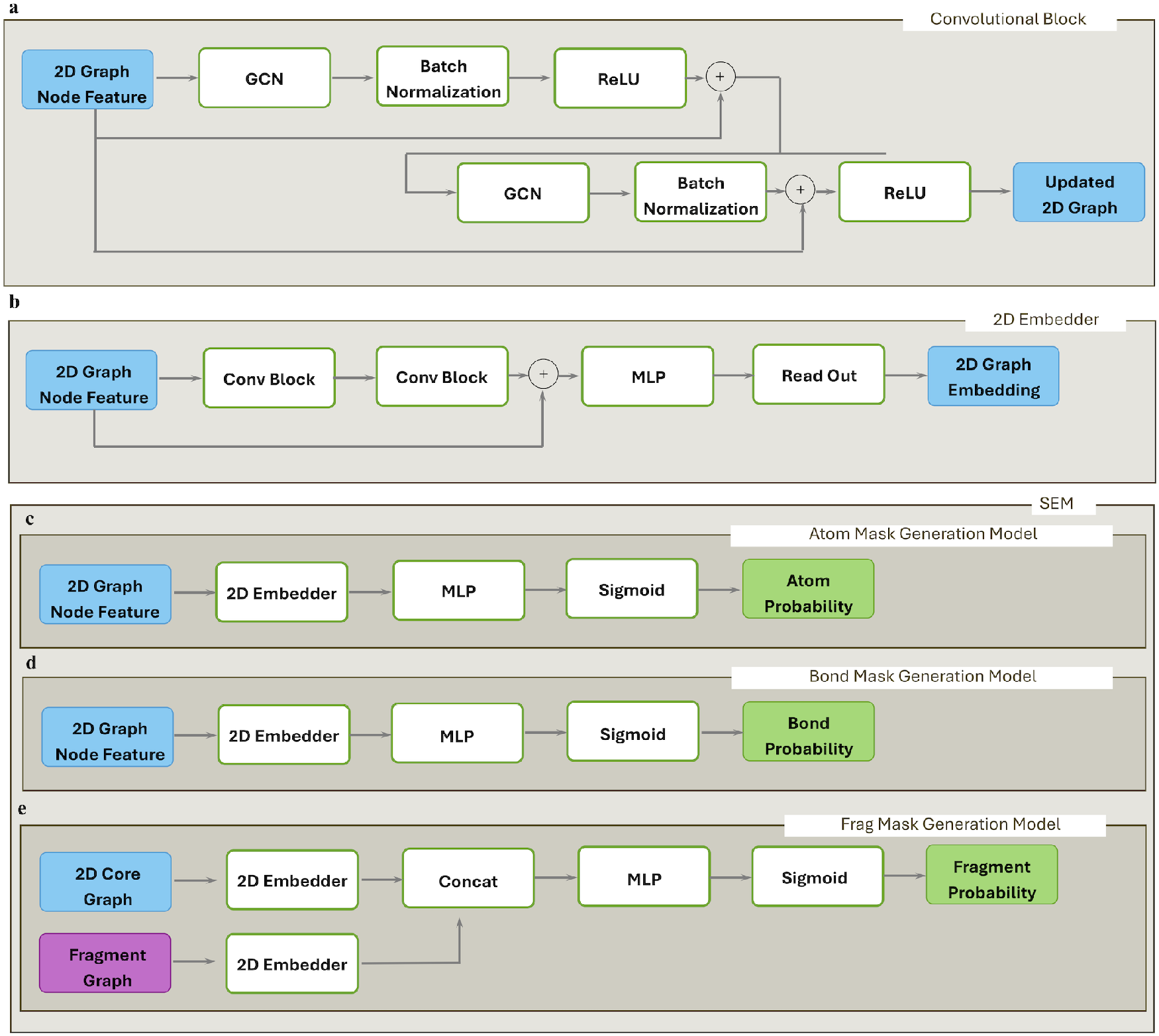
Detailed architecture of the SHARP deep learning model: (a) Convolutional block, (b) 2D embedder, (c–e) The SEM module, consisting of: (c) Atom mask generation model, (d) Bond mask generation model, (e) Fragment mask generation model.

**Figure 7:**
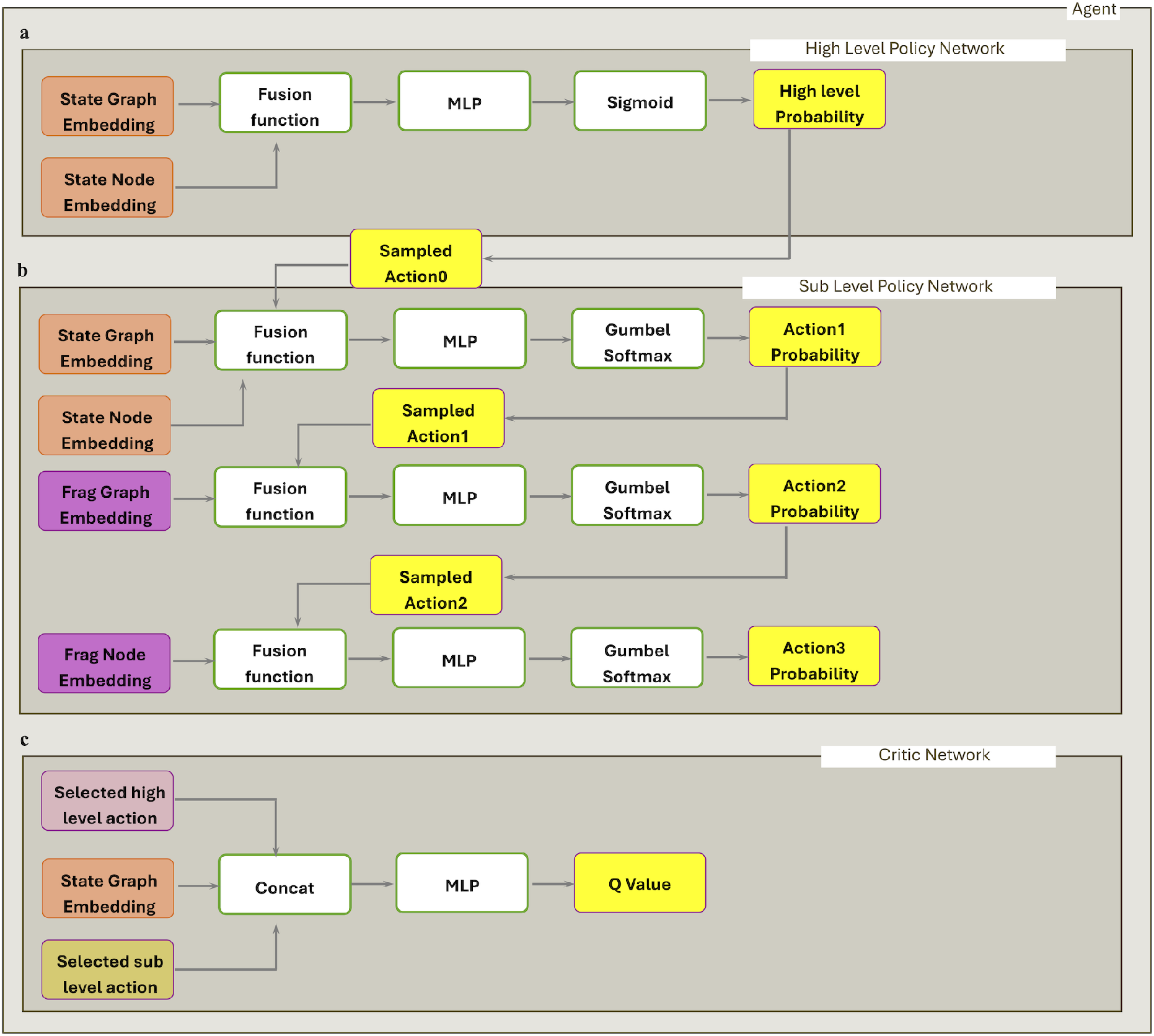
Detailed deep learning model architecture of SHARP. The agent consists of: (a) a high-level policy network, (b) a sub-policy network, and (c) a critic network.

#### The Synthesizability Estimation Model (SEM)

The model consists of three submodules: the Atom Mask Generation Model, Bond Mask Generation Model, and Fragment Mask Generation Model. The basic operation unit for the submodules is the Convolutional Block (Figure 6a) which consists of Graph Convolutional Network layers (GCNs).^37^ Then for each submodule, layers of Convolution blocks are used to build 2D embedder (Figure 6b) processing own input graph features to the next step.

The graphs used in SEM is generated by following steps. The workflow begins with a SMILES string. Using the RDKit library,^38^ this string is first parsed to create a complete molecule object, which understands the atoms and bonds involved. Next, an empty graph is initialized using the Deep Graph Library (DGL).^39^ The code then loops through every atom in the RDKit molecule, adding a corresponding node to the DGL graph. For each atom, a detailed feature vector is built to describe it numerically. This vector includes one-hot encoded representations of the atom symbol (from a vocabulary of C, N, O, S, P, F, H, Si, Cl, Br, I, and a wildcard *), the atom degree (number of neighbors, from 0 to 5), the total number of hydrogens (from 0 to 4), and the implicit valence (from 0 to 5). A final boolean flag (1 or 0) is added to this vector to specify if the atom is aromatic. This list of atom features is then converted into a PyTorch tensor and assigned to the graph’s nodes. Following the atoms, the program iterates through every chemical bond. For each bond, it adds edges to the DGL graph that connect the relevant nodes in both directions. A feature vector is then created for each bond, composed of several boolean flags. These flags identify the bond type (is it SINGLE, DOUBLE, TRIPLE, or AROMATIC?), whether the bond is conjugated, and if the bond is part of a ring. This collection of bond features is also converted into a PyTorch tensor and assigned to the graph’s edges. Finally, the completed DGL graph, now containing the full molecular structure and all the precisely defined atom and bond features, is returned as the output, ready for analysis.

For the Atom Mask Generation Model (Figure 6c) and Bond Mask Generation Model (Figure 6 d), 2D graph node features are taken as input, These features are passed through the 2D embedder, followed by a Multi-Layer Perceptron (MLP). Finally, a sigmoid activation function is applied to produce the atom mask or bond mask,

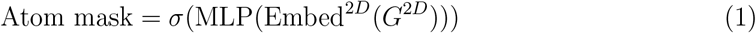

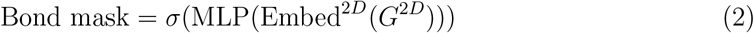

*h* indicates initial 2D graph’s node feature and *σ* indicates sigmoid function.

For the Frag Mask Generation Model (Figure 6 e), node features are brought from two graphs: 2D core graph and the fragment graph. These features are passed through two separate 2D embedders and are concatenated. Then, MLP and Sigmoid is passed and results in probability of those two fragments can be attached.

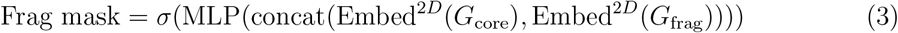

#### The Reinforcement Learning (RL) Agent

The RL agent consists of two components: a policy network and a critic network, as we employ the Soft Actor-Critic algorithm.^40^ Additionally, the policy network is divided into a high-level policy network and a sub-level policy network.

The process for generating the 3D graph of the ligand-protein complex begins by reading the atomic coordinates from a PDB file, which represents the docked structure. The code first identifies and separates the atoms belonging to the ligand from those of the protein. To focus on the immediate binding site, it creates a localized environment by selecting only the protein atoms that fall within a defined spatial box around the ligand’s center of mass. (25 Å) These selected protein atoms and all the ligand atoms together constitute the nodes of the graph. Each node is featurized with its 3D coordinates and a one-hot encoded vector representing its atomic element (e.g., C, N, O, S, P, F, H, SI, Cl, Br, I, *, unknown). The graph’s edges are established in two distinct ways: first, covalent bonds within the ligand are intelligently perceived using the OpenBabel library^41^ to ensure correct chemical structure; second, non-covalent interactions are inferred by creating edges between any ligand-protein atom pair whose distance is below a specific threshold. (4 Å) Finally, all this information—the nodes, the two types of edges, and their corresponding features—is compiled into a graph structure using the Deep Graph Library (DGL), creating a comprehensive representation of the binding interaction.

For all tasks, we utilize both the 3D protein-ligand complex graph *G*^3*D*^ and the 2D ligand graph *G*^2*D*^. The *G*^3*D*^ graph is embedded using an SE(3)-equivariant Graph Neural Network (GNN),^10^ while *G*^2*D*^ is embedded using the 2D embedder. The resulting embeddings are concatenated to form the graph embedding 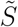 and the node embedding 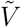. This process can be mathematically described as follows:

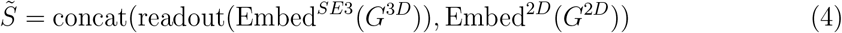

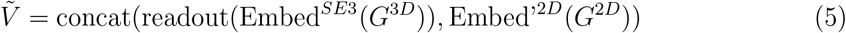

Embed’^2*D*^ indicates the 2D embedder without readout.

For the high-level policy, the inputs 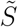 and 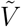 are fused through multiplicative interaction^42^ to pass the result through an MLP. (Figure 7a)The high-level policy probabilities *p*_0_ are then obtained using Gumbel-Softmax. ^43^ This process can be mathematically described as follows:

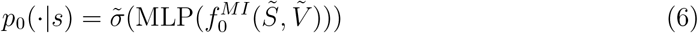

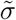 indicates Gumbel-Softmax,^43^ and 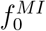 indicates multiplicative interaction. Then, *ã*_0_ is sampled from the probability.

We utilized the 4 actions as high level action: Fragment addition, Fragment deletion, and Fragment substitution. Each action needs to determine following sub-level policies:

- Fragment addition : i) which atom to add fragment in state molecule, ii) which fragment to add into state molecule, and iii) which attachment point to be used in fragment.
- Fragment deletion : i) which bond to break
- Fragment substitution : i) which bond to break, ii) which fragment to be add between them, and iii) how that fragment substituted

For each sub-level policy, sampling is performed at least three times autoregressively. (Figure 7 b) which do not affect the high-level policy. For actions such as fragment deletion, which requires only one step to complete, the process ends after sampling *ã*_1_, while other methods needs to sample *ã*_2_ and *ã*_3_.

The first step probability *p*_1_ to sample *a*_1_ is calculated by following equations:

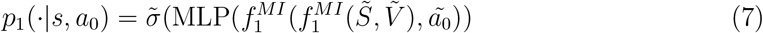

Before proceeding to the second step, which requires determining the fragment, all fragments needs to be embedded in the fragment library. This embedding process is carried out as follows:

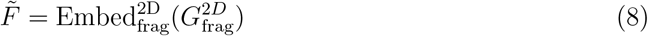

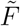 indicates concatenated embeddings of each fragment, and 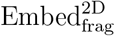 indicates independent 2D embedder for fragment.

Then by utilizing this, the probability *p*_2_ to sample *ã*_2_ is calculated as the following equation:

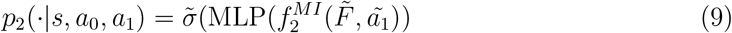

At third step, the probability *p*_3_ is calculated as follows:

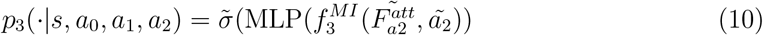

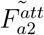 represents the node embedding of the attachment point from the embeddings of the fragment selected by *ã*_2_. Then, we sample the *ã*_3_.

For the critic network, the state information and all sampled actions are used as input to calculate the Q-value. (Figure 7 c) The process is mathematically represented by the following equation:

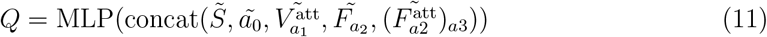

### Training Scheme

To train the atom mask generation model in SEM, fragments without attachment points were used as input, while the atom indices of the attachment points served as labels. Binary cross-entropy loss was applied to each atom index to determine whether it could serve as an attachment point. The loss function is as follows:

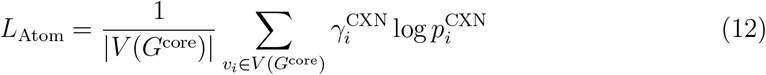

To train the bond mask generation model in SEM, the complete molecules from the original ChEMBL34 dataset were used as input, with the indices of BRICS breakable bonds serving as labels. Binary cross-entropy loss was applied to each bond index to predict whether it could be broken. Loss function is as follows:

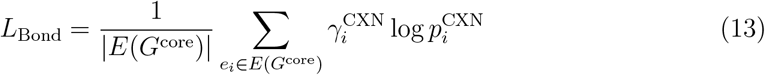

To train the fragment mask generation model in SEM, fragments originating from the same molecule were labeled as true, while fragments from different molecules were labeled as false, utilizing a negative sampling approach commonly applied in other studies. ^44^ Binary cross-entropy loss was used to predict whether a pair of fragments could bond together. The loss function is defined as follows:

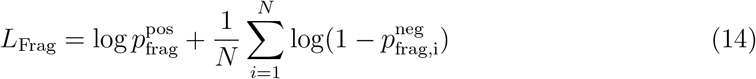

We utilized the Soft Actor-Critic(SAC) algorithm^40^ to train the model. The loss function to optimize the critic network is as follows.

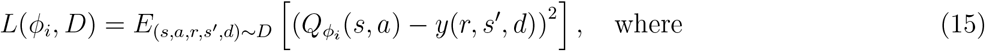

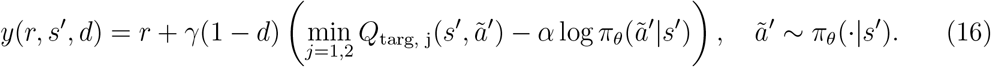

The loss function to optimize the policy network is as follows.

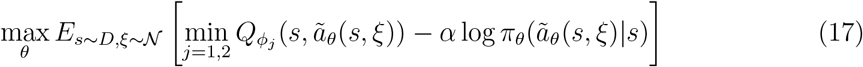

Hyperparameters were configured as follows: the embedder was set with an embedding dimension of 128 and comprised 3 layers. Training included 7 action steps per episode, with 3,000 exploration steps and a total of 5,000 steps. After the exploration phase, updates were performed every 50 steps, with 50 updates per interval. The optimizer was configured with a learning rate of 1 × 10^*−*4^, a batch size of 32 sampled from a replay buffer of size 1 × 10^6^, and the Adam optimizer.^45^

We handled the molecule, convert the file format and extracted chemical features using the RDKit,^38^ and Open Babel. ^41^ 3D docking poses and reward values are calculated using AutoDock Vina^46^ and AutoDock Vina-GPU.^47^ Given a pose, its protein-ligand interaction information is extracted using PLIP.^48^ Solvent accessible surface area is calculated using FreeSASA.^49^ Training and inference codes are written in Python3 using PyTorch^50^ and Scikit-learn^51^ libraries.

## Benchmark

### Quantitative performance benchmark

For the de novo design task, we generated 100 molecules equally across different methods (SHARP, Pocket2Mol,^12^ TargetDiff,^15^ GraphBP,^11^ DiffSBDD,^13^ LiGAN,^9^ and DiffBP^14^) for each target protein. To benchmark the performance of molecular generative models, we employ a comprehensive statistical analysis comparing various chemical properties. Our approach utilizes both traditional and novel metrics, with results typically presented in box plots and scatter plots.

Statistical significance between the models is assessed using two-sided tests: a Student’s t-test^52^ for normally distributed data and a Mann-Whitney U test^53^ for non-normally distributed data. Normality is evaluated using the Shapiro-Wilk test^54^ (for sample sizes less than 5000), while for larger samples, the t-test is applied due to the Central Limit Theorem. To account for multiple comparisons, a Bonferroni correction ^55^ is applied to the p-values in most analyses, ensuring robust statistical conclusions.

The benchmark incorporates several percentage-based metrics:

- Vina Score Percentage: Represents the proportion of generated molecules with a Vina score (indicating binding affinity) superior to protein-specific thresholds derived from our reference set. A higher percentage signifies better predicted binding affinity.
- Similarity Percentage: Indicates the proportion of generated molecules whose maximum fingerprint similarity to known high-binding-affinity (*k*_*i*_ *<* 1nM or IC_50_ *<* 1nM) molecules (from BindingDB or the test set, depending on recalculation) exceeds a protein-specific threshold. A higher percentage suggests generated molecules are chemically similar to known active compounds.
- SC Score Percentage: Quantifies the percentage of generated molecules with a Synthetic Complexity (SC) score below 4.0. A lower SC score indicates easier synthesizability, so a higher percentage here signifies greater synthetic accessibility.
- Unique Scaffold Percentage: Defined as the ratio of unique Bemis-Murcko scaffolds to the total number of generated molecules, expressed as a percentage. This metric reflects the internal diversity of the generated chemical space.
- Novel Scaffold Percentage: Represents the percentage of generated molecules whose Bemis-Murcko scaffolds are entirely novel (not present) within the reference set. A higher percentage indicates greater novelty and exploration of new chemical space by the model.

Additionally, Pareto fronts are calculated for pairs of objectives (e.g., Vina Score vs. Novelty) to visualize the trade-offs between different desired properties. SHARP’s Pareto front is specifically highlighted to facilitate direct comparison against other models, showcasing its ability to achieve optimal performance across multiple criteria simultaneously. Molecular diversity and novelty are further visualized using t-distributed Stochastic Neighbor Embedding (t-SNE) plots of molecular fingerprints, which provide a low-dimensional representation of the chemical space explored by each model in relation to the training and test sets.

### 3D interaction fingerprint analysis

To characterize protein-ligand interactions, we employed the PLIP tool^48^ to extract detailed interaction information from 3D complex structures. For each protein-ligand complex, PLIP identifies various interaction types, including hydrogen bonds, salt bridges, hydrophobic contacts, pi-stacking, pi-cation interactions, halogen bonds, water bridges, and metal complexes. We converted these interactions into a binary fingerprint vector. Each position in the vector corresponds to a specific residue number and interaction type. A value of ‘1’ indicates the presence of that interaction, while ‘0’ indicates its absence. This approach allows for a quantitative comparison of interaction profiles between different molecules.

#### Diversity and novelty analysis based on interaction fingerprints

We analyzed the generated interaction fingerprints to assess both the diversity and novelty of the generated molecules.

- Novelty: For each generated molecule, we calculated its Jaccard distance (1.0 - Jaccard similarity^56^) to the most similar molecule in the reference set. This distance represents the novelty of the interaction pattern, with higher values indicating greater dissimilarity from known interactions.
- Diversity: We calculated the pairwise Jaccard similarity between all generated molecules. To quantify diversity, we performed single-linkage clustering on these pairwise distances, using distance thresholds 0.6.

#### t-SNE visualization of interaction fingerprints

To visualize the high-dimensional interaction fingerprint data, we used t-distributed Stochastic Neighbor Embedding (t-SNE). t-SNE reduces the dimensionality of the interaction fingerprints to two dimensions, allowing us to plot the molecules in a 2D space where similar interaction patterns are clustered together. We used the scikit-learn implementation of t-SNE, with a perplexity parameter of 30 (or the maximum possible value if the number of samples was less than 31) and a random state of 42 for reproducibility. We generated t-SNE plots both for all molecules combined and for molecules interacting with individual proteins. In these plots, density contours, generated using Kernel Density Estimation, highlight the distribution of different groups (generated molecules, reference set, and test set).

### Learning and action analysis

We conducted a detailed analysis of the molecular generation trajectories to understand the model’s behavior during optimization. The trajectory data, which includes molecular SMILES strings and corresponding docking rewards at each step, was loaded and processed. Molecular properties (number of rings, atoms, and molecular weight) were also calculated for each generated molecule.

Our statistical analysis focused on several key aspects of the generation process:

- Reward Progression: To evaluate the learning progress, we plotted the median docking reward at each step, with the shaded region representing the standard deviation. The statistical significance of the reward increase over steps was quantified using the Pearson correlation coefficient between the step number and the docking reward.
- Action Distribution: We analyzed the frequency of different action types (addition, deletion, substitution) across distinct phases of the molecular generation. Specifically, we compared the action distribution in early steps (1-4) versus later steps (7-10) of the optimization process. The statistical difference in these distributions was assessed using a Chi-square test of independence, and the effect size was quantified by Cramer’s V.
- Fragment Substitution Characteristics: For fragment substitution actions, we examined the correlation between the properties (number of atoms and molecular weight) of the original and substituted fragments. This was done using the Pearson correlation coefficient. A high correlation would suggest that the model prefers substitutions with fragments of similar sizes, indicating fine-tuning of the molecular structure rather than drastic changes. The statistical significance of these correlations was also reported.
- Fragment Diversity: We analyzed the diversity of fragments utilized by the model by counting the occurrences of each unique fragment. The Shannon entropy was calculated to quantify the overall diversity of fragment usage. Additionally, a histogram of fragment usage frequency, a bar chart of the top 20 most frequently used fragments, and a cumulative distribution plot of fragment usage were generated to visually represent the diversity. The fragment distribution of generated molecules was also compared with that of the training and test sets using Jaccard similarity and the Kolmogorov-Smirnov (KS) two-sample test on their normalized frequency distributions. This comparison assesses the model’s ability to generate novel fragments while maintaining relevance to the target chemical space.

### Rationality evaluation on generated molecules

To assess the physicochemical properties and drug-likeness of generated molecules, we computed several molecular descriptors for compounds produced by each generative model. These descriptors include atom count, rotatable bonds, hydrogen bond donors (HBD), hydrogen bond acceptors (HBA), molecular weight (MW), and LogP. For comparative analysis, we also included a curated subset of molecules from the ChEMBL34 database as a reference (size : 2,409,270).

For each molecular property, we generated box plots to visualize the distribution of values across different models and the ChEMBL34 reference set. Statistical significance in property distributions between models and against the ChEMBL34 reference was determined using a two-sided statistical test. The choice of test depended on the data’s normality: for sample sizes under 5000, we performed the Shapiro-Wilk test for normality; if the data were normally distributed (*p >* 0.05), a Student’s t-test with unequal variances was applied. For non-normally distributed data, or for sample sizes of 5000 or greater (where the Central Limit Theorem generally allows for parametric tests), the Mann-Whitney U test was used. To account for multiple comparisons, a Bonferroni correction was applied to all reported p-values.

To evaluate the drug-likeness of generated compounds, we calculated the percentage of molecules from each model that comply with Lipinski’s Rule of Five. This rule defines drug-like molecules as those meeting the following criteria: Molecular Weight ≤ 500 Da, LogP ≤ 5, Hydrogen Bond Donors ≤ 5, and Hydrogen Bond Acceptors ≤ 10.

In addition to standard molecular properties, we specifically analyzed the distribution of ring sizes (3-membered, 4-membered, 7-membered, and *>*7-membered rings) within the generated molecular sets. For each ring size, we calculated the percentage of generated molecules containing at least one such ring. These percentages were then visualized using separate bar plots for each ring size category.

Box plots displayed the distributions of molecular properties, with each model represented by a distinct color, including black for SHARP and gray for ChEMBL34. Percentages reflecting Lipinski’s rule compliance or specific ring statistics were annotated directly above the corresponding bars. Statistical significance was indicated with p-value annotations above the box plots, linking each model’s distribution to the reference distribution (ChEMBL34). Legends for all plots were generated separately to maintain visual clarity and conciseness within the main figures.

### Lead optimization tasks

For the scaffold hopping task, we generated 100 molecules for each target protein and each scaffold in the high-binding-affinity list. Similar to the de novo design task, we identified the most similar molecule from the generated set of 100 molecules to the high-binding-affinity list for the corresponding receptor. From each scaffold for each protein, we selected one representative molecule from the generated set and compiled a list. Finally, we calculated the mean similarity across this list. Validity is calculated over all the attempts to generate the molecules.

For the remaining tasks, we generated 100 molecules for each target protein and each input fragment from the high-binding-affinity list. Among the 100 generated molecules, we identified the most recovered molecule based on fingerprint similarity. The similarity was measured for each receptor and fragment, and the mean similarity was calculated across all cases. Validity was evaluated in the same manner as in the scaffold hopping task.

To assess our model’s lead optimization capabilities on diverse datasets, we utilized data from the literature on fragment-to-lead optimization and quantitative structure-activity relationship (QSAR) studies.^57^ We initially evaluated 8 biological systems that met the following criteria: (i) availability of 3D structures of the initial fragment-protein complex in the Protein Data Bank (PDB),^58^ (ii) not being a protein-protein interaction (PPI) system, and (iii) not covalent bond inhibitors as reported by Woodhead et al.. Although this source primarily focuses on fragment-to-lead optimization, some systems involved substantial initial fragment modification, allowing us to categorize them as scaffold hopping tasks. Conversely, systems with minimal initial fragment modification were categorized as sidechain decoration tasks. As no explicit linker design tasks were available, we adapted one system by fragmenting a molecule with terminal rings to create a linker design problem. From these, we selected four systems that yielded particularly impressive results for our case studies. Protein-ligand interaction figures were generated using Maestro,^59^ with an interaction cutoff set at 4 Å. For each task and test set molecule, we performed 10 generation steps and produced 100 molecules.

## Data availability

All datasets used in SHARP, including the training, validation, and benchmark sets, are available at https://figshare.com/articles/dataset/SHARP_Generating_Synthesizable_Molecules_via_Fragment-based_Hierarchical_Action-space_Reinforcement_Learning_for_Pareto_Optimization/29597885. The data were collected from the open-source Cross-Docked2020 dataset^32^ and BindingDB.^35^

## Code availability

The SHARP code repository, including the model architecture, training scripts, inference pipeline, and analysis tools, is available at: https://github.com/jasonkim8652/SHARP

## Acknowledgements

This work was supported by the National Research Foundation of Korea (NRF) grant (RS-2024-00407331 to CS), Institute of Information & communications Technology Planning & Evaluation (IITP) grant (RS-2023-00220628 to CS), Korea-US Collaborative Research Fund(KUCRF) grant (RS-2024-00467483 to CS and HP), and National Research Foundation of Korea (NRF) grant (RS-2024-00407331 to CS, 2022R1C1C1007817 to HP) funded by the Korea government (MIST).

## Author contributions

Jeonghyeon Kim, Seongok Ryu, Hahnbeom Park, and Chaok Seok contributed to the conception of the idea, implementation, and experiments. Jeonghyeon Kim drafted the manuscript, and all authors—Jeonghyeon Kim, Seongok Ryu, Hahnbeom Park, and Chaok Seok—participated in manuscript revision.

## Supplementary Information

### t-SNE analysis for each protein

We conducted t-SNE analyses for each protein system using both 2D structural fingerprints (Figure 8) and 3D protein–ligand interaction fingerprints (Figure 9). In both analyses, across all protein targets, molecules generated by SHARP consistently occupied a broader area in the t-SNE space compared to those from other methods. Moreover, SHARP’s molecular distribution was consistently located farther from the reference set and closer to the test set, suggesting a superior ability to explore novel and relevant regions of chemical space.

**Figure 8:**
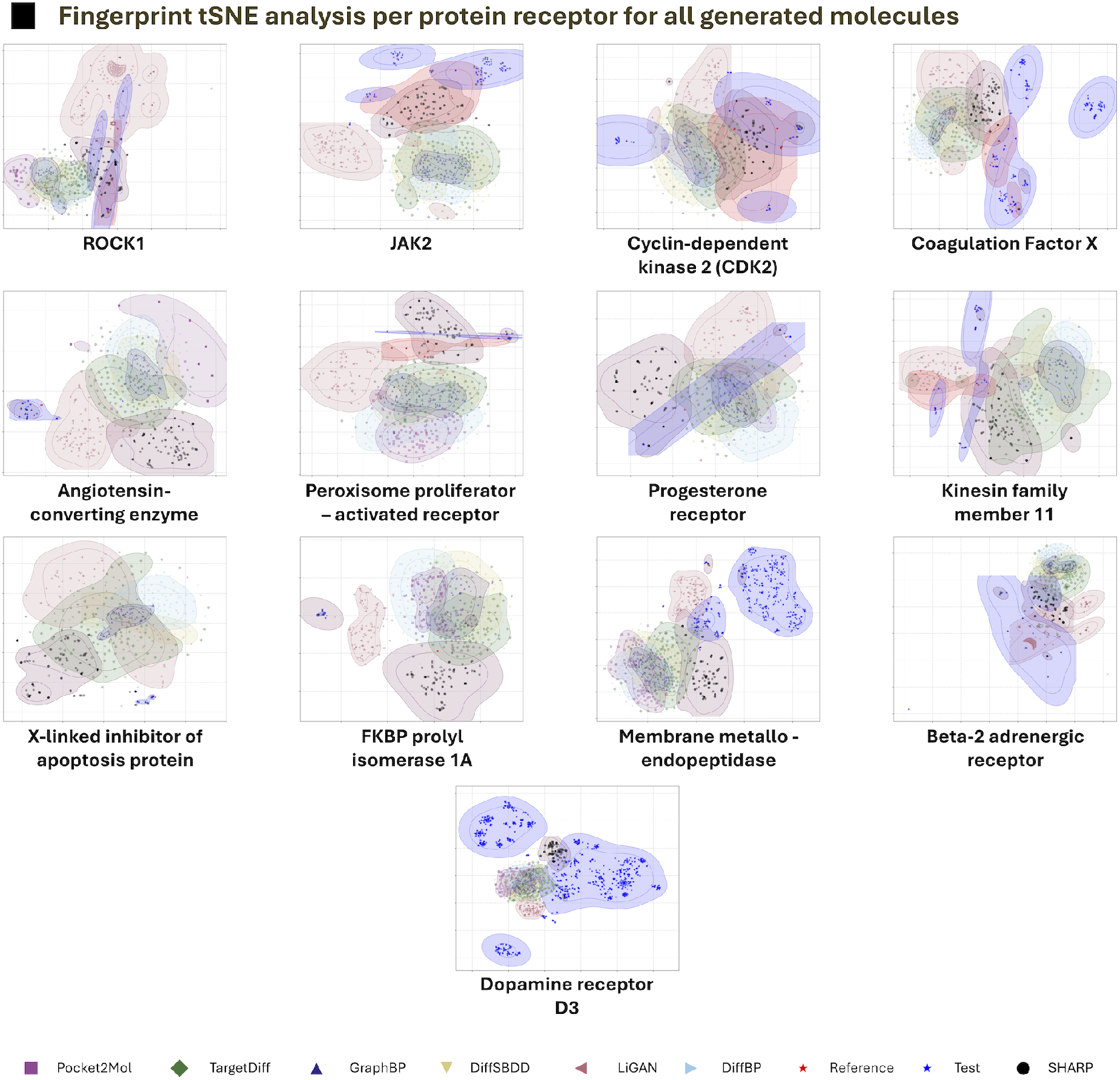
t-SNE visualization of 2D structural fingerprints for each protein system. Molecules generated by different models are color-coded, with the legend provided at the bottom of the figure.

**Figure 9:**
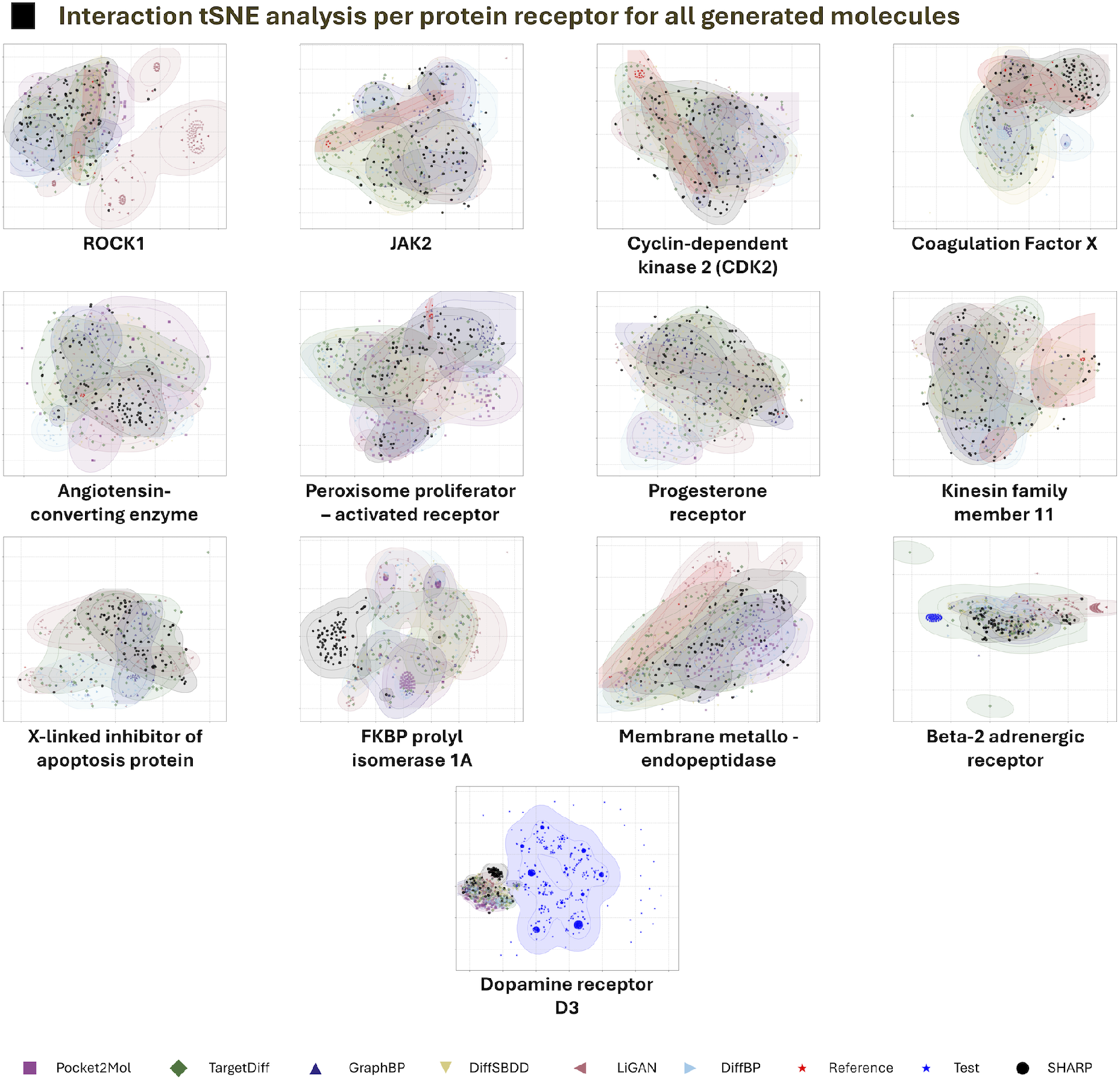
t-SNE visualization of 3D protein–ligand interaction fingerprints for each protein system. Molecules generated by different models are color-coded, with the corresponding legend shown at the bottom of the figure.

### Binding pose analysis of generated molecules

**Figure 10:**
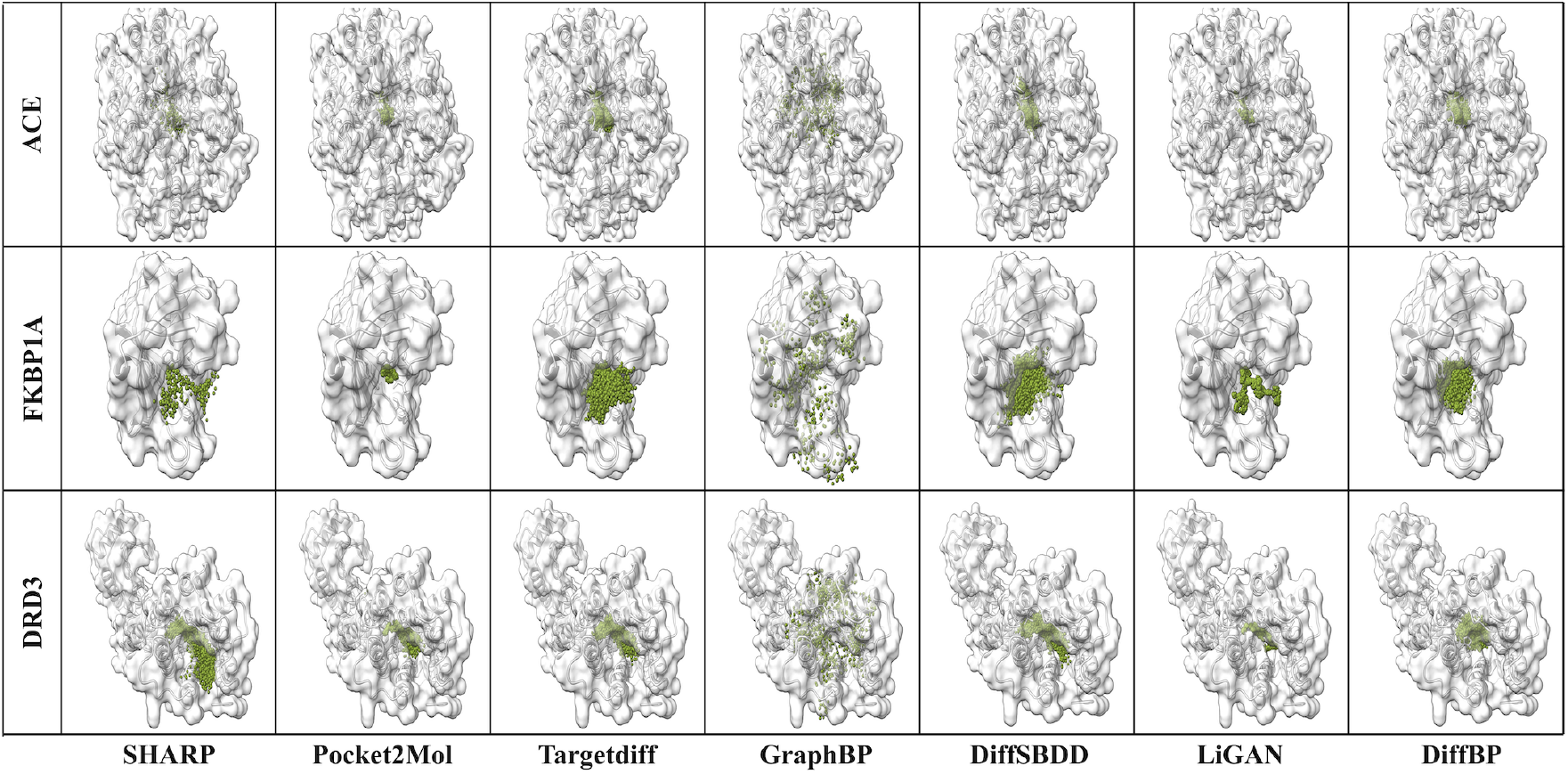
Binding pose analysis of generated molecules in De novo drug design task for receptors Angiotensin-converting enzyme(ACE), FKBP1A, and Dopamine D3 receptor(DRD3). Green points represent each generated molecule’s atomic positions.

We evaluated whether the generated molecules were positioned rationally within the binding pocket. To do this, receptor-contacting atoms of each generated molecule were highlighted in green, as shown in Figure 4a. We selected three receptors from distinct protein families for this analysis: angiotensin-converting enzyme (ACE), FKBP1A, and the dopamine D3 receptor (DRD3). Among them, ACE possesses a more enclosed binding pocket, while FKBP1A and DRD3 feature relatively open binding sites.

The diversity and quality of generated molecular positions vary significantly across different generation methods. GraphBP produces molecules with a highly sparse and dispersed distribution of atomic positions, regardless of whether the binding pocket is enclosed or open. This often leads to steric clashes with protein atoms, and in some cases, atoms are placed far from the binding site. In contrast, LiGAN and Pocket2Mol do not exhibit problematic atomic placements; however, their positional coverage is relatively narrow. These methods generate molecules by conditioning on pharmacophore information from existing data, which improves the hit rate but limits molecular diversity (Table 1).

Other methods show no major issues in atomic positioning and achieve better diversity. Notably, in proteins with open binding sites, such as FKBP1A and DRD3, SHARP tends to place atoms toward more exposed regions of the pocket, demonstrating broader spatial exploration compared to other approaches. This wider exploration is a key advantage in de novo drug design, as it increases structural diversity, improves the likelihood of discovering novel hit compounds, and enhances understanding of the binding pocket’s landscape.

### Examples of generated molecules for each design task

In this section, we present a case study to evaluate the effectiveness of each model on four tasks: fragment growing, linker design, scaffold hopping, and sidechain decoration. To assess performance, we visualized the binding modes of generated molecules for each task using different target receptors from diverse families. For the fragment growing task, we selected the β2-adrenergic receptor (ADRB2) as the target protein (Figure 5a). For linker design, we used cyclin-dependent kinase 2 (CDK2) (Figure 5b). Kinesin-like protein 1 (KIF11) was chosen for the scaffold hopping task (Figure 5c), and the progesterone receptor (PRGR) for sidechain decoration (Figure 5d). In all tasks, DiffSBDD frequently produced invalid molecular structures with abnormal atomic positions, as illustrated in Figure 5. As a result, we exclude DiffSBDD from further analysis in this section.

For the fragment growing task, SHARP, Pocket2Mol, and TargetDiff produced broadly similar results. GraphBP, however, added an excessive number of atoms from the outset, hindering its ability to achieve an optimal fit within the binding pocket. TargetDiff generated the most minimal and efficient extension, while Pocket2Mol produced a nearly correct solution. Notably, SHARP was the only model to exactly reproduce the original ligand structure, highlighting its superior accuracy in this task.

For the linker design task, both SHARP and TargetDiff generated accurate and comparable results. In contrast, GraphBP failed to properly recognize both fragments, adding atoms only to one side and placing them outside the binding pocket—suggesting a poor un derstanding of the pocket’s geometry. Pocket2Mol formed a highly strained four-membered ring, likely due to its strategy of focusing primarily on generating a linker with reasonable bond lengths, without adequately considering geometric and energetic constraints. TargetDiff demonstrated a robust approach by successfully designing a linker that satisfied the spatial and structural constraints between the two fragments. SHARP also produced a correct and well-placed linker, further highlighting its accuracy in this task.

**Figure 11:**
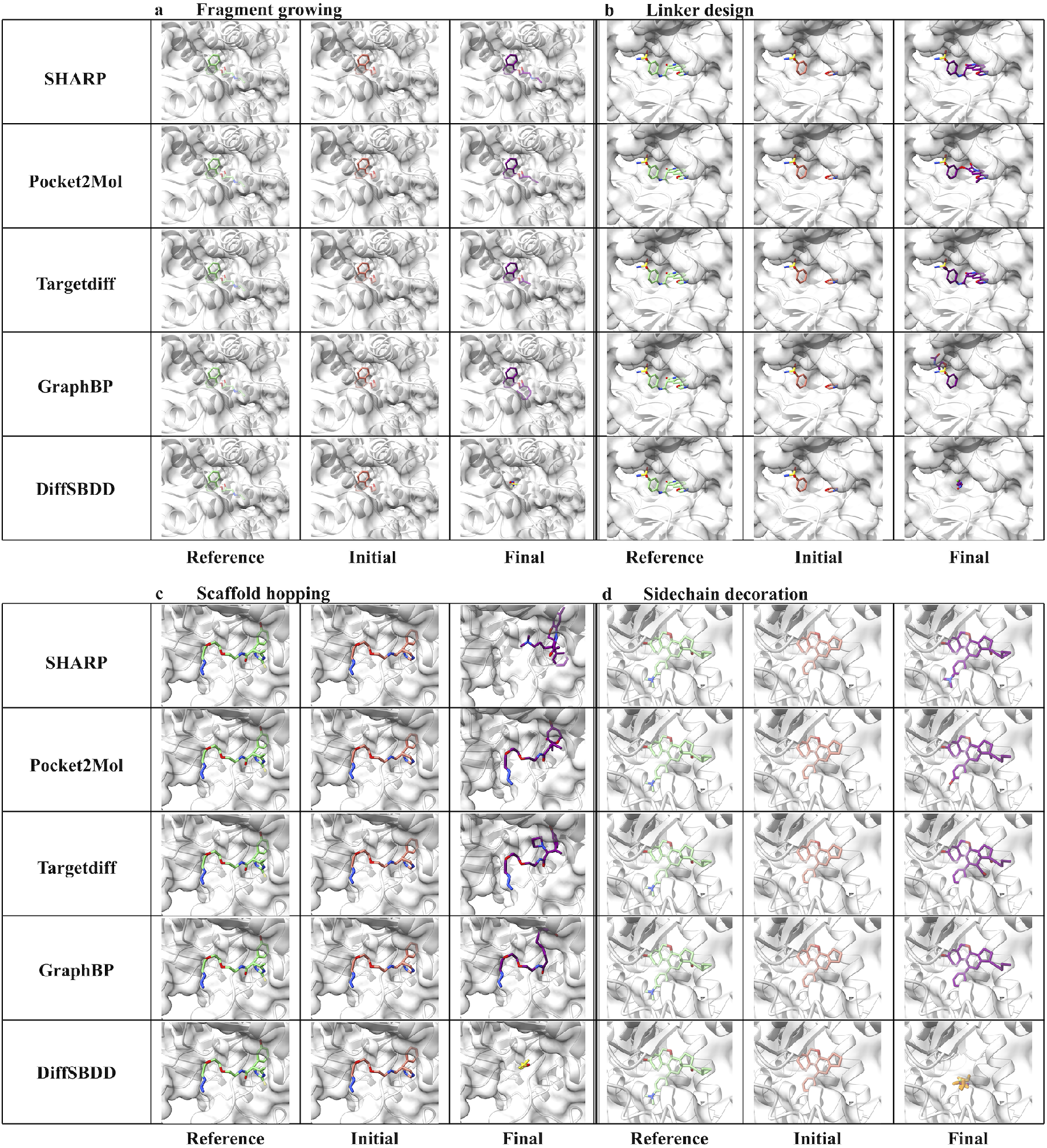
Examples of molecules generated by each model for the following tasks: (a) Fragment growing with the β2-adrenergic receptor (ADRB2), (b) Linker design with cyclin-dependent kinase 2 (CDK2), (c) Scaffold hopping with kinesin-like protein 1 (KIF11), and (d) Sidechain decoration with the progesterone receptor (PRGR).

For the scaffold hopping task, all methods except SHARP made only minor modifications to the input scaffold, despite clear opportunities for improvement from a medicinal chemistry perspective—for example, the presence of too many rotatable bonds. GraphBP exacerbated this issue by further elongating the rotatable bonds, resulting in molecules likely to exhibit low binding affinity. TargetDiff and Pocket2Mol also failed to address this structural limitation, with TargetDiff additionally generating a strained four-membered ring. In contrast, SHARP successfully identified the limitations of the original scaffold and explored alternative interactions deeper within the binding pocket. Remarkably, the new scaffold proposed by SHARP matched known high-affinity ligands, underscoring its ability to redesign scaffolds effectively. These results highlight SHARP’s strength in scaffold hopping: recognizing structural weaknesses in the original molecule, discovering new interaction sites, and generating novel scaffolds distinct from the initial input.

For the sidechain decoration task, GraphBP made the fewest modifications to the input structure, despite the clear need for more extensive changes based on the reference ligand. TargetDiff, in contrast, modified irrelevant regions while overlooking critical areas, indicating a limited understanding of the sidechain’s role in optimizing binding affinity. Pocket2Mol targeted the appropriate regions but did not sufficiently modify them to match the reference. Notably, SHARP made all necessary modifications and produced an exact match to the reference ligand, demonstrating superior accuracy and task-specific understanding in sidechain decoration.

## References

(1) Bohacek, R. S.; McMartin, C.; Guida, W. C. The art and practice of structure-based drug design: a molecular modeling perspective. Medicinal research reviews 1996, 16, 3–50.

(2) Xiong, G.; Wu, Z.; Yi, J.; Fu, L.; Yang, Z.; Hsieh, C.; Yin, M.; Zeng, X.; Wu, C.; Lu, A.; others ADMETlab 2.0: an integrated online platform for accurate and comprehensive predictions of ADMET properties. Nucleic acids research 2021, 49, W5–W14.

(3) Cumming, J. G.; Davis, A. M.; Muresan, S.; Haeberlein, M.; Chen, H. Chemical predictive modelling to improve compound quality. Nature reviews Drug discovery 2013, 12, 948–962.

(4) Hou, T.; Wang, J. Structure–ADME relationship: still a long way to go? Expert Opinion on Drug Metabolism & Toxicology 2008, 4, 759–770.

(5) Loeffler, H. H.; He, J.; Tibo, A.; Janet, J. P.; Voronov, A.; Mervin, L. H.; Engkvist, O. Reinvent 4: Modern AI–driven generative molecule design. Journal of Cheminformatics 2024, 16, 20.

(6) Kang, S.; Cho, K. Conditional molecular design with deep generative models. Journal of chemical information and modeling 2018, 59, 43–52.

(7) Jumper, J.; Evans, R.; Pritzel, A.; Green, T.; Figurnov, M.; Ronneberger, O.; Tunyasuvunakool, K.; Bates, R.; Žídek, A.; Potapenko, A.; others Highly accurate protein structure prediction with AlphaFold. nature 2021, 596, 583–589.

(8) Baek, M.; DiMaio, F.; Anishchenko, I.; Dauparas, J.; Ovchinnikov, S.; Lee, G. R.; Wang, J.; Cong, Q.; Kinch, L. N.; Schaeffer, R. D.; others Accurate prediction of protein structures and interactions using a three-track neural network. Science 2021, 373, 871–876.

(9) Ragoza, M.; Masuda, T.; Koes, D. R. Generating 3D molecules conditional on receptor binding sites with deep generative models. Chemical science 2022, 13, 2701–2713.

(10) Du, W.; Zhang, H.; Du, Y.; Meng, Q.; Chen, W.; Zheng, N.; Shao, B.; Liu, T.-Y. SE (3) equivariant graph neural networks with complete local frames. International Conference on Machine Learning. 2022; pp 5583–5608.

(11) Liu, M.; Luo, Y.; Uchino, K.; Maruhashi, K.; Ji, S. Generating 3d molecules for target protein binding. arXiv preprint 2204.09410 2022,

(12) Peng, X.; Luo, S.; Guan, J.; Xie, Q.; Peng, J.; Ma, J. Pocket2mol: Efficient molecular sampling based on 3d protein pockets. International Conference on Machine Learning. 2022; pp 17644–17655.

(13) Schneuing, A.; Harris, C.; Du, Y.; Didi, K.; Jamasb, A.; Igashov, I.; Du, W.; Gomes, C.; Blundell, T.; Lio, P.; others Structure-based drug design with equivariant diffusion models. arXiv preprint 2210.13695 2022,

(14) Lin, H.; Huang, Y.; Zhang, O.; Ma, S.; Liu, M.; Li, X.; Wu, L.; Wang, J.; Hou, T.; Li, S. Z. Diffbp: Generative diffusion of 3d molecules for target protein binding. arXiv preprint 2211.11214 2022,

(15) Guan, J.; Qian, W. W.; Peng, X.; Su, Y.; Peng, J.; Ma, J. 3d equivariant diffusion for target-aware molecule generation and affinity prediction. arXiv preprint 2303.03543 2023,

(16) Gao, W.; Luo, S.; Coley, C. W. Generative Artificial Intelligence for Navigating Synthesizable Chemical Space. arXiv preprint 2410.03494 2024,

(17) Ertl, P.; Schuffenhauer, A. Estimation of synthetic accessibility score of drug-like molecules based on molecular complexity and fragment contributions. Journal of cheminformatics 2009, 1, 1–11.

(18) Pateria, S.; Subagdja, B.; Tan, A.-h.; Quek, C. Hierarchical reinforcement learning: A comprehensive survey. ACM Computing Surveys (CSUR) 2021, 54, 1–35.

(19) Xie, W.; Zhang, J.; Xie, Q.; Gong, C.; Ren, Y.; Xie, J.; Sun, Q.; Xu, Y.; Lai, L.; Pei, J. Accelerating discovery of bioactive ligands with pharmacophore-informed generative models. Nature communications 2025, 16, 2391.

(20) Bemis, G. W.; Murcko, M. A. The properties of known drugs. 1. Molecular frameworks. Journal of medicinal chemistry 1996, 39, 2887–2893.

(21) Gaulton, A.; Bellis, L. J.; Bento, A. P.; Chambers, J.; Davies, M.; Hersey, A.; Light, Y.; McGlinchey, S.; Michalovich, D.; Al-Lazikani, B.; others ChEMBL: a large-scale bioactivity database for drug discovery. Nucleic acids research 2012, 40, D1100–D1107.

(22) Pollastri, M. P. Overview on the Rule of Five. Current protocols in pharmacology 2010, 49, 9–12.

(23) Böhm, H.-J.; Flohr, A.; Stahl, M. Scaffold hopping. Drug discovery today: Technologies 2004, 1, 217–224.

(24) Hu, Y.; Stumpfe, D.; Bajorath, J. Recent advances in scaffold hopping: miniperspective. Journal of medicinal chemistry 2017, 60, 1238–1246.

(25) Smith, C. R.; Aranda, R.; Bobinski, T. P.; Briere, D. M.; Burns, A. C.; Christensen, J. G.; Clarine, J.; Engstrom, L. D.; Gunn, R. J.; Ivetac, A.; others Fragment-based discovery of MRTX1719, a synthetic lethal inhibitor of the PRMT5• MTA complex for the treatment of MTAP-deleted cancers. Journal of Medicinal Chemistry 2022, 65, 1749–1766.

(26) Charoensutthivarakul, S.; Thomas, S. E.; Curran, A.; Brown, K. P.; Belardinelli, J. M.; Whitehouse, A. J.; Acebrón-García-de Eulate, M.; Sangan, J.; Gramani, S. G.; Jackson, M.; others Development of inhibitors of SAICAR synthetase (PurC) from Mycobacterium abscessus using a fragment-based approach. ACS infectious diseases 2022, 8, 296–309.

(27) Luttens, A.; Gullberg, H.; Abdurakhmanov, E.; Vo, D. D.; Akaberi, D.; Talibov, V. O.; Nekhotiaeva, N.; Vangeel, L.; De Jonghe, S.; Jochmans, D.; others Ultralarge virtual screening identifies SARS-CoV-2 main protease inhibitors with broad-spectrum activity against coronaviruses. Journal of the American Chemical Society 2022, 144, 2905–2920.

(28) Mandal, M.; Xiao, L.; Pan, W.; Scapin, G.; Li, G.; Tang, H.; Yang, S.-W.; Pan, J.; Root, Y.; de Jesus, R. K.; others Rapid evolution of a fragment-like molecule to pan-metallo-beta-lactamase inhibitors: initial leads toward clinical candidates. Journal of Medicinal Chemistry 2022, 65, 16234–16251.

(29) Jeon, W.; Kim, D. Autonomous molecule generation using reinforcement learning and docking to develop potential novel inhibitors. Scientific reports 2020, 10, 22104.

(30) Korshunova, M.; Huang, N.; Capuzzi, S.; Radchenko, D. S.; Savych, O.; Moroz, Y. S.; Wells, C. I.; Willson, T. M.; Tropsha, A.; Isayev, O. Generative and reinforcement learning approaches for the automated de novo design of bioactive compounds. Communications Chemistry 2022, 5, 129.

(31) Chen, B.; Fu, X.; Barzilay, R.; Jaakkola, T. Fragment-based sequential translation for molecular optimization. arXiv preprint 2111.01009 2021,

(32) Francoeur, P. G.; Masuda, T.; Sunseri, J.; Jia, A.; Iovanisci, R. B.; Snyder, I.; Koes, D. R. Three-dimensional convolutional neural networks and a cross-docked data set for structure-based drug design. Journal of chemical information and modeling 2020, 60, 4200–4215.

(33) Sankar, S.; Chandran Sakthivel, N.; Chandra, N. Fast local alignment of protein pockets (FLAPP): a system-compiled program for large-scale binding site alignment. Journal of Chemical Information and Modeling 2022, 62, 4810–4819.

(34) Mysinger, M. M.; Carchia, M.; Irwin, J. J.; Shoichet, B. K. Directory of useful decoys, enhanced (DUD-E): better ligands and decoys for better benchmarking. Journal of medicinal chemistry 2012, 55, 6582–6594.

(35) Gilson, M. K.; Liu, T.; Baitaluk, M.; Nicola, G.; Hwang, L.; Chong, J. BindingDB in 2015: a public database for medicinal chemistry, computational chemistry and systems pharmacology. Nucleic acids research 2016, 44, D1045–D1053.

(36) Degen, J.; Wegscheid-Gerlach, C.; Zaliani, A.; Rarey, M. On the art of compiling and using’drug-like’chemical fragment spaces. ChemMedChem 2008, 3, 1503.

(37) Kipf, T. N.; Welling, M. Semi-supervised classification with graph convolutional networks. arXiv preprint 1609.02907 2016,

(38) Bento, A. P.; Hersey, A.; Félix, E.; Landrum, G.; Gaulton, A.; Atkinson, F.; Bellis, L. J.; De Veij, M.; Leach, A. R. An open source chemical structure curation pipeline using RDKit. Journal of Cheminformatics 2020, 12, 1–16.

(39) Wang, M. Y. Deep graph library: Towards efficient and scalable deep learning on graphs. ICLR workshop on representation learning on graphs and manifolds. 2019.

(40) Haarnoja, T.; Zhou, A.; Abbeel, P.; Levine, S. Soft actor-critic: Off-policy maximum entropy deep reinforcement learning with a stochastic actor. International conference on machine learning. 2018; pp 1861–1870.

(41) O’Boyle, N. M.; Banck, M.; James, C. A.; Morley, C.; Vandermeersch, T.; Hutchison, G. R. Open Babel: An open chemical toolbox. Journal of cheminformatics 2011, 3, 1–14.

(42) Jayakumar, S. M.; Czarnecki, W. M.; Menick, J.; Schwarz, J.; Rae, J.; Osindero, S.; Teh, Y. W.; Harley, T.; Pascanu, R. Multiplicative interactions and where to find them. International conference on learning representations. 2020.

(43) Jang, E.; Gu, S.; Poole, B. Categorical reparameterization with gumbel-softmax. arXiv preprint 1611.01144 2016,

(44) Robinson, J.; Chuang, C.-Y.; Sra, S.; Jegelka, S. Contrastive learning with hard negative samples. arXiv preprint 2010.04592 2020,

(45) Kingma, D. P. Adam: A method for stochastic optimization. arXiv preprint 1412.6980 2014,

(46) Trott, O.; Olson, A. J. AutoDock Vina: improving the speed and accuracy of docking with a new scoring function, efficient optimization, and multithreading. Journal of computational chemistry 2010, 31, 455–461.

(47) Ding, J.; Tang, S.; Mei, Z.; Wang, L.; Huang, Q.; Hu, H.; Ling, M.; Wu, J. Vina-GPU 2.0: further accelerating AutoDock Vina and its derivatives with graphics processing units. Journal of chemical information and modeling 2023, 63, 1982–1998.

(48) Salentin, S.; Schreiber, S.; Haupt, V. J.; Adasme, M. F.; Schroeder, M. PLIP: fully automated protein–ligand interaction profiler. Nucleic acids research 2015, 43, W443–W447.

(49) Mitternacht, S. FreeSASA: An open source C library for solvent accessible surface area calculations. F1000Research 2016, 5.

(50) Paszke, A.; Gross, S.; Massa, F.; Lerer, A.; Bradbury, J.; Chanan, G.; Killeen, T.; Lin, Z.; Gimelshein, N.; Antiga, L.; others Pytorch: An imperative style, high-performance deep learning library. Advances in neural information processing systems 2019, 32.

(51) Pedregosa, F.; Varoquaux, G.; Gramfort, A.; Michel, V.; Thirion, B.; Grisel, O.; Blondel, M.; Prettenhofer, P.; Weiss, R.; Dubourg, V.; others Scikit-learn: Machine learning in Python. the Journal of machine Learning research 2011, 12, 2825–2830.

(52) Student The probable error of a mean. Biometrika 1908, 6, 1–25.

(53) Mann, H. B.; Whitney, D. R. On a test of whether one of two random variables is stochastically larger than the other. The annals of mathematical statistics 1947, 18, 50–60.

(54) Shapiro, S. S.; Wilk, M. B. An analysis of variance test for normality (complete samples). Biometrika 1965, 52, 591–611.

(55) Dunn, O. J. Multiple comparisons among means. Journal of the American Statistical Association 1961, 56, 52–64.

(56) Ivchenko, G.; Honov, S. On the jaccard similarity test. Journal of Mathematical Sciences 1998, 88, 789–794.

(57) Woodhead, A. J.; Erlanson, D. A.; de Esch, I. J.; Holvey, R. S.; Jahnke, W.; Pathuri, P. Fragment-to-lead medicinal chemistry publications in 2022. Journal of medicinal chemistry 2024, 67, 2287–2304.

(58) Burley, S. K.; Berman, H. M.; Kleywegt, G. J.; Markley, J. L.; Nakamura, H.; Velankar, S. Protein Data Bank (PDB): the single global macromolecular structure archive. Protein crystallography: methods and protocols 2017, 627–641.

(59) Maestro, S. Maestro. Schrödinger, LLC, New York, NY 2020, 2020, 682.

